# Differential polysaccharide utilization is the basis for a nanohaloarchaeon : haloarchaeon symbiosis

**DOI:** 10.1101/794461

**Authors:** Violetta La Cono, Enzo Messina, Manfred Rohde, Erika Arcadi, Sergio Ciordia, Francesca Crisafi, Renata Denaro, Manuel Ferrer, Laura Giuliano, Peter N. Golyshin, Olga V. Golyshina, John E. Hallsworth, Gina La Spada, Maria C. Mena, Margarita A. Shevchenko, Francesco Smedile, Dimitry Y. Sorokin, Stepan V. Toshchakov, Arcady Mushegian, Michail M. Yakimov

## Abstract

Nanohaloarchaeota, a clade of diminutive archaea, with small genomes and limited metabolic capabilities, are ubiquitous in hypersaline habitats, which they share with the extremely halophilic and phylogenetically distant euryarchaea. Some of these nanohaloarchaeota and euryarchaea appear to interact with each other. In this study, we investigate the genetic and physiological nature of their relationship. We isolated the nanohaloarchaeon *Candidatus* Nanohalobium constans LC1Nh and the haloarchaeon *Halomicrobium* sp. LC1Hm from a solar saltern, reproducibly co-cultivated these species, sequenced their genomes, and characterized their metabolic/trophic interactions. The nanohaloarchaeon is a magnesium-dependent aerotolerant heterotrophic anaerobe of the DPANN superphylum; it lacks respiratory complexes and its energy production relies on fermentative metabolism of sugar derivatives, obtained by depolymerizing alpha-glucans or by acquiring the chitin monomer N-acetylglucosamine from the chitinolytic haloarchaeal host. *Halomicrobium* is a member of the class *Halobacteria* and a chitinotrophic aerobe. The nanohaloarchaeon lacks key biosynthetic pathways and is likely to be provided with amino acids, lipids, nucleotides and cofactors via physical contact with its host *Halomicrobium*. In turn, the ability of *Ca*. Nanohalobium to hydrolyse alpha-glucans boosts the host’s growth in the absence of a chitin substrate. These findings suggest that at least some members of the nanohaloarchaea, previously considered ecologically unimportant given their limited metabolic potential, in fact may play significant roles in the microbial carbon turnover, food chains, and ecosystem function. The behaviour of *Halomicrobium*, which accommodates the colonization by *Ca*. Nanohalobium, can be interpreted as a bet-hedging strategy, maximizing its long-term fitness in a habitat where the availability of carbon substrates can vary both spatially and temporarily.

## Introduction

Hypersaline lakes and solar salterns with salt concentrations close to the saturation point house unique consortia of extremely halophilic organisms. Despite harsh physico-chemical conditions in these habitats, communities of halophiles are dense and diverse, comprising representatives of all domains of life, i.e., extremely halophilic prokaryotes (Bacteria and Archaea), unicellular green algae, protists and arthropods (brine shrimps and in some cases also brine flies)^1^. These arthropods have chitinous exoskeletons and reach considerable biomass (up to 50 g m^-2^), making chitin one of the most abundant biopolymers in these ecosystems^2-4^. Halophilic fermentative bacteria are known to play a role in chitin mineralization in hypersaline habitats worldwide^5-7^, and more recently it was shown that haloarchaea also take part in the primary degradation of polymeric organic matter in many of the same habitats^8-11^. Here we reveal the role of haloarchaea as chitinotrophic hosts for nanohaloarchaea, the extremely halophilic members of the DPANN superphylum^12-15^.

Members of phylum “*Ca*. Nanohaloarchaeota” were first detected less than a decade ago, in 0.22 µm-filtered samples collected from Spanish solar salterns and the Australian hypersaline Lake Tyrell^16,17^. Recent environmental surveys of 16S rRNA amplicons, metagenomic sequences and lineage-specific quantitative FISH of cells from natural samples indicate that nanohaloarchaea thrive in hypersaline ecosystems worldwide, often representing a notable fraction of the total archaeal community^17-20^. To date, few finished genome sequences are available for the representatives of DPANN superphylum, and difficulties in their cultivation prevented proper characterization of their growth requirements, morphology and physiology. Only ectosymbiotic and ectoparasitic thermophiles from the candidate phylum Nanoarchaeota and acidophiles from the candidate phylum Micrarchaeota have been obtained in stable binary co-cultures with their hosts from the phyla Crenarchaeota and Euryachaeota, respectively^21-24^. Most recently, the study of enrichment cultures containing members of nanohaloarchaea (*Ca*. Nanoarchaeum antarcticus), and other microbial strains including heterotrophic haloarchaeon *Halorubrum lacusprofundi*, was published by Hamm *et al*. (2019)^25^. This exciting development supports the hypothesis of Andrade *et al*. (2015)^19^ that some haloarchaea may act as hosts for nanohaloarchaea.

In this study, we report the enrichment, cultivation and characterization of a binary association of the nanohaloarchaeon LC1Nh, dubbed ‘*Ca*. Nanohalobium constans’ (latter on – *Ca*. Nanohalobium), and its host, a chitinotrophic euryarchaeon *Halomicrobium* sp. LC1Hm, from the hypersaline marine solar saltern Saline della Laguna (Trapani, Italy). We used a finished ungapped genome sequence of the nanohaloarchaeon LC1Nh in conjunction with the electron microscopy, growth analysis, proteomic and metabolomic data, to uncover and dissect the mechanisms of mutually beneficial interactions between these archaeal associates and for the first time reveal the ecological importance of nanohaloarchaea.

## Results

Surface sediments and near-bottom brine from the crystallizer pond of the artisanal family-run salt work *Saline della Laguna* located in western Sicily (Italy) were used to enrich for chitinotrophic haloarchaea (Supplementary Fig. 1). A pinkish-coloured culture was obtained after 16 weeks in mineral medium LC (Laguna Chitin) (salinity 240 g l^-1^, 3.42 M Na^+^ and 0.32 M Mg^2+^) by microaerophilic incubation (without shaking) in the dark at 37°C. The medium was supplemented with bacteria-suppressing antibiotics vancomycin and streptomycin (100 mg l^-1^ each) and shrimp chitin (5 g l^-1^) as a single source of energy and carbon. After two 1:100 passag’s, taxonomic analysis of 16S rRNA gene sequences from the enriched chitinotrophic community revealed a 10-fold increase of *Halomicrobium* and or *Halomicrobium*-related haloarchaea, compared to their initial presence in the crystallizer pond, accompanied by a similar increase in OTUs representing putative nanohaloarchaea (Supplementary Fig. 2). Following three rounds of dilution-to-extinction transfers with the same antibiotic selection, the enrichment was examined microscopically and analysed by sequencing of amplified 16S rRNA genes. A binary culture of nanohaloarchaeal and haloarchaeal species was obtained. The haloarchaeon (strain LC1Hm) was closely related to the representatives of the genus *Halomicrobium*, which includes two members known to have chitinolytic activities^9^. Based on the at the nucleotide sequence similarity level of 99.3-99.8% to other members of *Halomicrobium*, this isolate can be assigned to the genus. The 16S rRNA tree, constructed from the alignment of the 16S rRNA gene of the putative nanohaloarchaeon (strain LC1Nh) to the master alignment in SILVA Release 132SSURef NR99^26^, and inferred using maximum parsimony criteria within the ARB software^27^, suggests that the nearest neighbour of the novel nanohaloarchaeon was the partially sequenced Cry7_clone28 (GenBank GQ374969, 94.3% identity) from Dry Creek crystallizer pond (Australia) and other partially sequenced clones (≈91% identity) from around the world, all belonging to Clade III of the candidate phylum *Nanohaloarchaeota* (Supplementary Fig. 3). According to the rRNA phylogeny, the LC1Nh nanohaloarchaeon is distinct from the other candidate genera in the phylum (86.5 – 91.8% sequence identities) and falls within the range of recently recommended values (86.5 and 94.5%) for the family- and genus-level classifications^28^. This was confirmed by a phylogenomic analysis of a concatenated alignment of the 122 single-copy archaeal marker proteins using the Genome Taxonomy Data Base (GTDB) framework^29^. The *Nanohaloarchaeota* phylum appears to be split into two class-level taxa, each with some cultivated representatives, as well as a deep “*Ca*. Nanosalinarium” cluster of uncertain taxonomic rank (Fig. 1; Supplementary Fig. 4). We propose a new class “*Ca.* Nanohalobia” that includes a new order, “*Ca*. Nanohalobiales” and the novel family, “*Ca*. Nanohalobiaceae”, which in turn accommodates the genus “*Ca*. Nanohalobium”.

**Figure 1.**
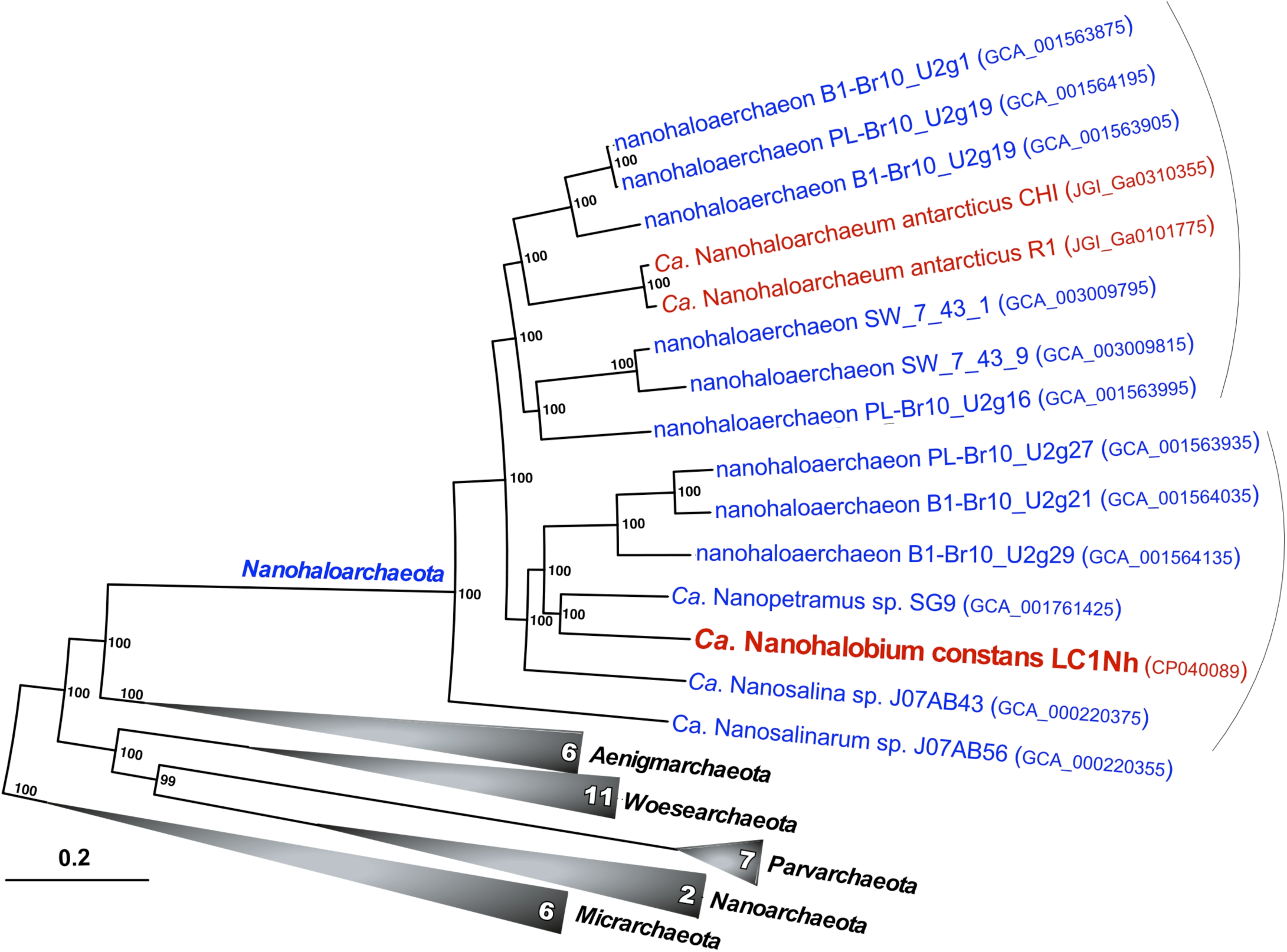
Phylogenetic placement of *Ca*. Nanohalobium constans LC1Nh based on concatenated partial amino acid sequences of the 122 proteins conserved in Archaea (archaeal marker genes). Bootstrap values are shown at the nodes. Bar, 0.20 changes per position. Cultivated and uncultured members of candidate phylum *Nanohaloarchaeota* are highlighted in red and blue, respectively. Detailed list of genomes and methods used for the tree construction is given in Materials and Methods.

The LC1Nh nanohaloarchaeon was observed in the co-culture as small coccoid cells either free-living or associated with large pleomorphic host cells (Fig. 2; Supplementary Fig. 1). Transmission (TEM) and field emission scanning electron microscopic (FESEM) imaging revealed a regular S-layer-like outer surface of the nanohaloarchaeal cells (Figs. 2d-i). The cocci formed an association with almost 20% of the cells within the *Halomicrobium* population, at the median multiplicity of 4-5, but occasionally reaching 17 cells per host cell (Figs. 2d-f; Supplementary Fig. 1c). In some cases, a single nanoarchaeal cell bridged two host cells (Figs. 2g,h). When grown on chitin, *Halomicrobium* sp. LC1Hm cells produced a thick, electron-dense external layer, which often engulfed the symbiont (Fig. 3). This layer was not observed during growth on monosaccharides (N-acetylglucosamine [GlcNAc] and dextrose) or disaccharides (maltose and cellobiose) (Supplementary Fig. 5), suggesting that this extracellular material is involved in the interaction of the cells with the polysaccharide particles, similarly to what was recently demonstrated for cellulolytic natronoarchaea of the genus *Natronobifoma*^30^.

**Figure 2.**
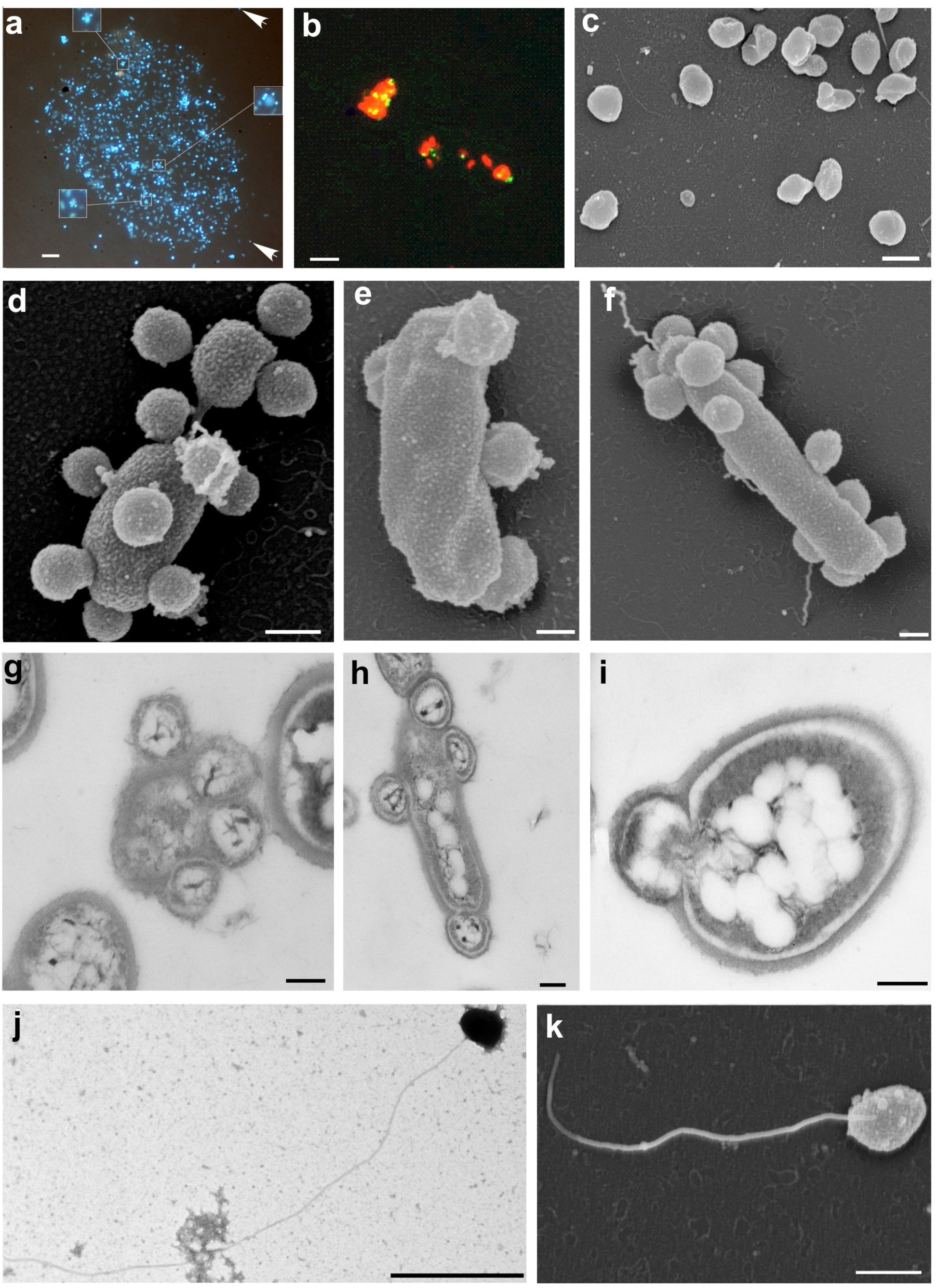
Fluorescence, TEM and FESEM micrographs of nanohaloarchaeote *Ca*. Nanohalobium constans LC1Nh and its chitinotrophic host *Halomicrobium* sp. LC1Hm cells. DAPI (**a**) and CARD-FISH (**b**) staining shows tiny coccoidal nanohaloarchaeal cells (285 ± 50 nm in diameter) either detached or adhering to the host haloarchaeal cells contact with host haloarchaea (tyramide Alexa488 and Alexa594 were used for nanohaloarchaeal and haloarchaeal cells, respectively); FESEM image depicts the coccoidal shape of detached nanohaloarchaeota cells (**c**); up to 17 nanohaloarchaeota cells can closely interact with the host *Halomicrobium* sp. LC1Hm cell (**d-f**); some nanohaloarchaeal cells can be attached to more than one host (**g**,**h**); a series of TEM ultrathin sections depict the intimate contact of nanohaloarchaeota cells with its chitinolytic host resulting in the formation of membrane depression (stretching) on the host cell surface (**g-i**); nanohaloarchaeal cells express long archaella (**j**,**k**). Bars represent 5 µm in (**a-b**), 0.2 µm in (**c-I, k**) and 0.5 µm in (**j**).

**Figure 3.**
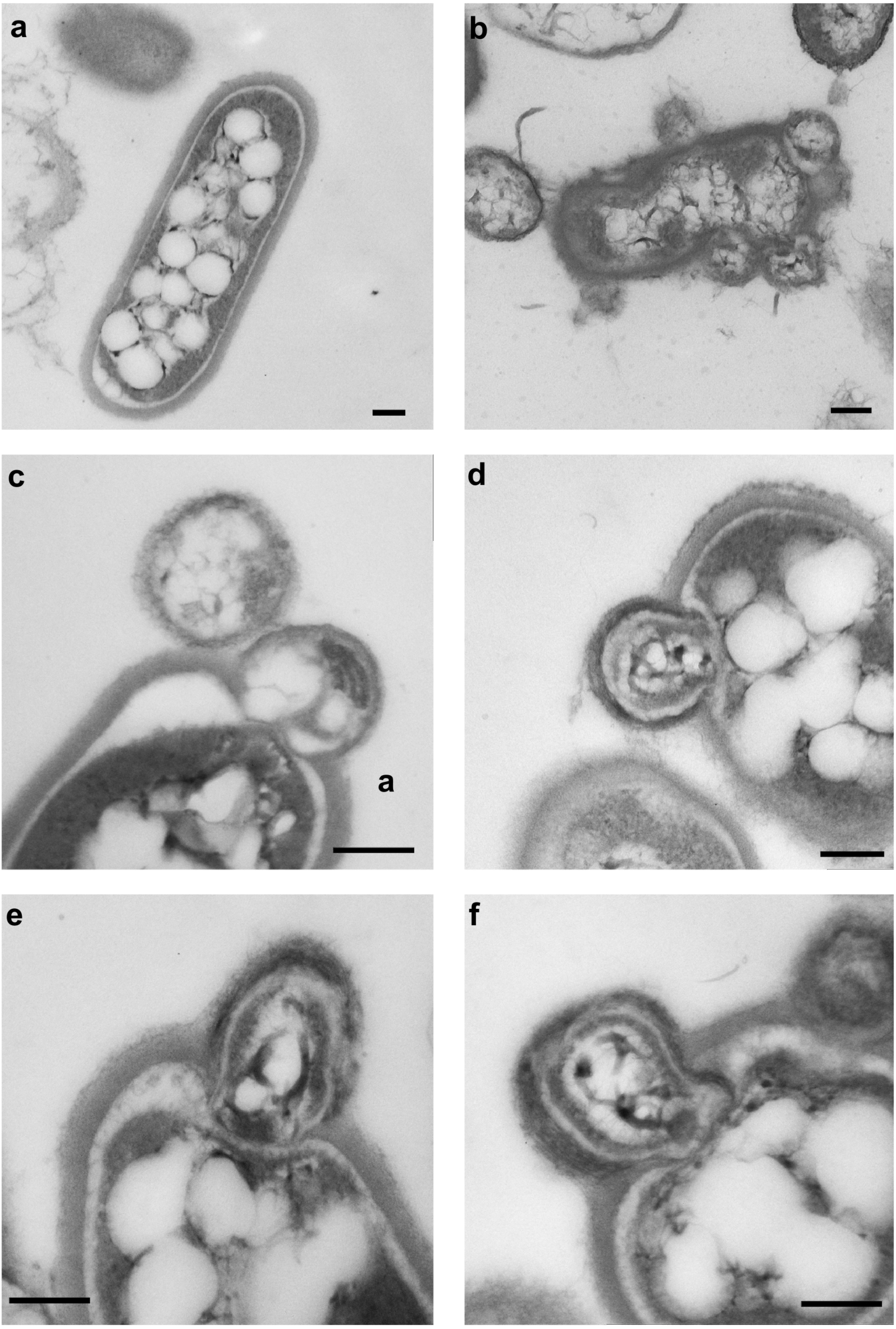
TEM microscopy of chitin-growing co-culture of *Halomicrobium* sp. LC1Hm and *Ca*. Nanohalobium constans LC1Nh. (**a**) symbiont-free *Halomicrobium* sp. LC1Hm cell; (**b**) pronounced membrane stretching and (**c**) exfoliation of a thick electron-dense external layer of the host cell; (c-f) various stages of cells fusion suggestive of a shared lipid membrane and extracellular matrix. Bars represent 500 nm.

Thin-section TEM imaging of both chitin- and GlcNAc-grown LC1Nh-LC1Hm co-cultures revealed various aspects of interaction between the nanohaloarchaeon and the host. The initial surface attachment with visibly separated boundary layer was frequently accompanied by stretching of the host membrane. At some micrographs there were clear signs of nanohaloarchaeon penetration through the host S-layer and even occasional breaching of the host cell membrane (Fig. 3). The predatory-like behaviour may be required for the nanohaloarchaeon in order to acquire biosynthetic precursors and metabolites from the host. This is consistent with the analysis of gene repertoire in *Ca*. Nanohalobium, which apparently lacks key biosynthetic pathways for amino acids, lipids, nucleotides and cofactors (see below). In addition, TEM and FESEM micrographs of nanohaloarchaeal cells showed pili-like structures made of thick and long monotrichous or ditrichous protein stalks, frequently unwinding to thin filaments (Figs. 2j,k; Supplementary Figs. 1c-e). It remains to be seen whether these structures are used for motility of detached nanohaloarcheal cells, or to perform an alternative role, such as attachment to host cells or to polysaccharide substrate.

To separate the chitinotrophic host from the nanohaloarchaeon, we plated aliquots of the co-culture on solid LC medium supplemented with amorphous chitin as a sole substrate. The developed colonies of axenic *Halomicrobium* sp. LC1Hm culture formed large clearance zones, indicative of the extracellular production of chitin hydrolases. The unattached nanohaloarchaeal cells were separated from the haloarchaeal cells by three rounds of filtering the LC1Hm+LC1Nh co-culture through a 0.45-µm membrane filter. Unlike *Halomicrobium* sp. LC1Hm, which could be maintained in pure culture, the nanohaloarchaeon did not grow alone, even when provided with different energy sources under various cultivation conditions, including the addition of the supernatant from *Halomicrobium* sp. LC1Hm culture or lysates of its biomass. These cultivation experiments, in which the nutrients from the growing *Halomicrobium* sp. LC1Hm culture were provided while physical contact of the nanohaloarchaeon and the host was prevented, suggest that cell–cell contact is required for LC1Nh to proliferate. In contrast, mixing the membrane-filtered nanohaloarchaeal cell suspension with an axenic culture of *Halomicrobium* sp. LC1Hm gave rise to a reconstructed co-culture, suggesting that at least a fraction of the LC1Nh nanohaloarchaeon remained viable in the free state and capable of re-establishing the attachment to the host. Thus, the presence of haloarchaeal host appears to be mandatory for cultivation of nanohaloarchaeal ectosymbiont under given conditions, but not *vice versa*. Mixing the isolated *Ca*. Nanohalobium cells with pure cultures of other polysaccharidolytic archaea, such as chitinolytic *Natrinema* sp. HArcht2 and *Salinarchaeum* sp. HArcht-Bsk1^9^ or amylolytic *Halorhabdus thiamatea*^31^, as well as sugar-utilizing haloarchaeon *Haloferax volcanii* DSM3757, failed to produce a stable co-culture. Indeed, the nanohaloarchaeon was completely eliminated from these mixtures after less than two weeks of incubation. These findings suggest that the nanohaloarchaeon depends upon a specific interaction with *Halomicrobium* sp. LC1Hm. We named this new nanohaloarchaeon ‘*Candidatus* Nanohalobium constans*’* (the nanosized salt-loving organism, constant in its allegiance to a specific host).

In the axenic culture of the host, and in the host-ectosymbiont co-culture, chitin was completely degraded after less than two weeks of incubation at 240 g l^-1^ salinity and 40°C under microaerophilic conditions (initial dissolved oxygen at <4.0 mg l^-1^). The presence of the ectosymbiont in a co-culture slightly prolonged the lag phase of the host, though neither the yield of the host biomass nor its maximum specific growth rates (µ_max_ = 0.028-0.035 h^-1^) were affected (Fig. 4a). Only N-acetylglucosamine (GlcNAc), the monomeric product of chitin degradation, was found in either of the supernatants, but none of the low molecular weight chitodextrin oligomers, (GlcNAc)_2-6_, were detected. Complete degradation of insoluble chitin to GlcNAc usually occurs via two types of hydrolytic enzymes: endochitinases, which digest chitin microfibrils at internal sites to form chitodextrins and N,N’-diacetylchitobiose, and exochitodextrinases, which further break down chitodextrin oligomers at their termini to produce monosaccharide GlcNAc. All this suggests that chitodextrin oligomers, except perhaps those longer than six sugar residues, are short-lived and are rapidly converted to monomeric product in the co-culture. It is noteworthy that GlcNAc was detected in both supernatants at relatively high concentrations (3.7 ± 0.9 and 5.2 ± 1.6 mmol in axenic and ectosymbiont-host cultures, respectively). Other characteristics of *Halomicrobium* behaviour also differed in the presence and absence of the ectosymbiont. For example, the growth of host-ectosymbiont association on chitin was accompanied by threefold higher accumulation of acetate (6.39 ± 1.55 vs. 2.10 ± 0.33 mmol, *P* = 0.002) compared with axenic host culture (Fig. 4b). The host-ectosymbiont association also consumed twice as much oxygen as the pure *Halomicrobium* culture (*P* ≤ 0.009), resulting in microaerobic conditions at the early stationary phase of growth (Fig. 4c; see Supplementary Text for more detail). In a separate experiment, we established that *Halomicrobium* sp. LC1Hm is an obligate aerobe, whereas the nanohaloarchaeon did not reproduce under the well-oxygenated regime optimal for the host (>4 mg l^-1^ of dissolved oxygen). Thus, only microaerobic conditions facilitate the growth of both organisms in a co-culture. Besides the requirement for a low oxygen tension, another parameter was also mandatory for the prosperity of the host-ectosymbiont co-culture. This was the presence of significant amount of magnesium; the host-ectosymbiont co-culture can be stably maintained only in the range from 75 to 800 mM of Mg^2+^ (Supplementary Fig. 6).

**Figure 4.**
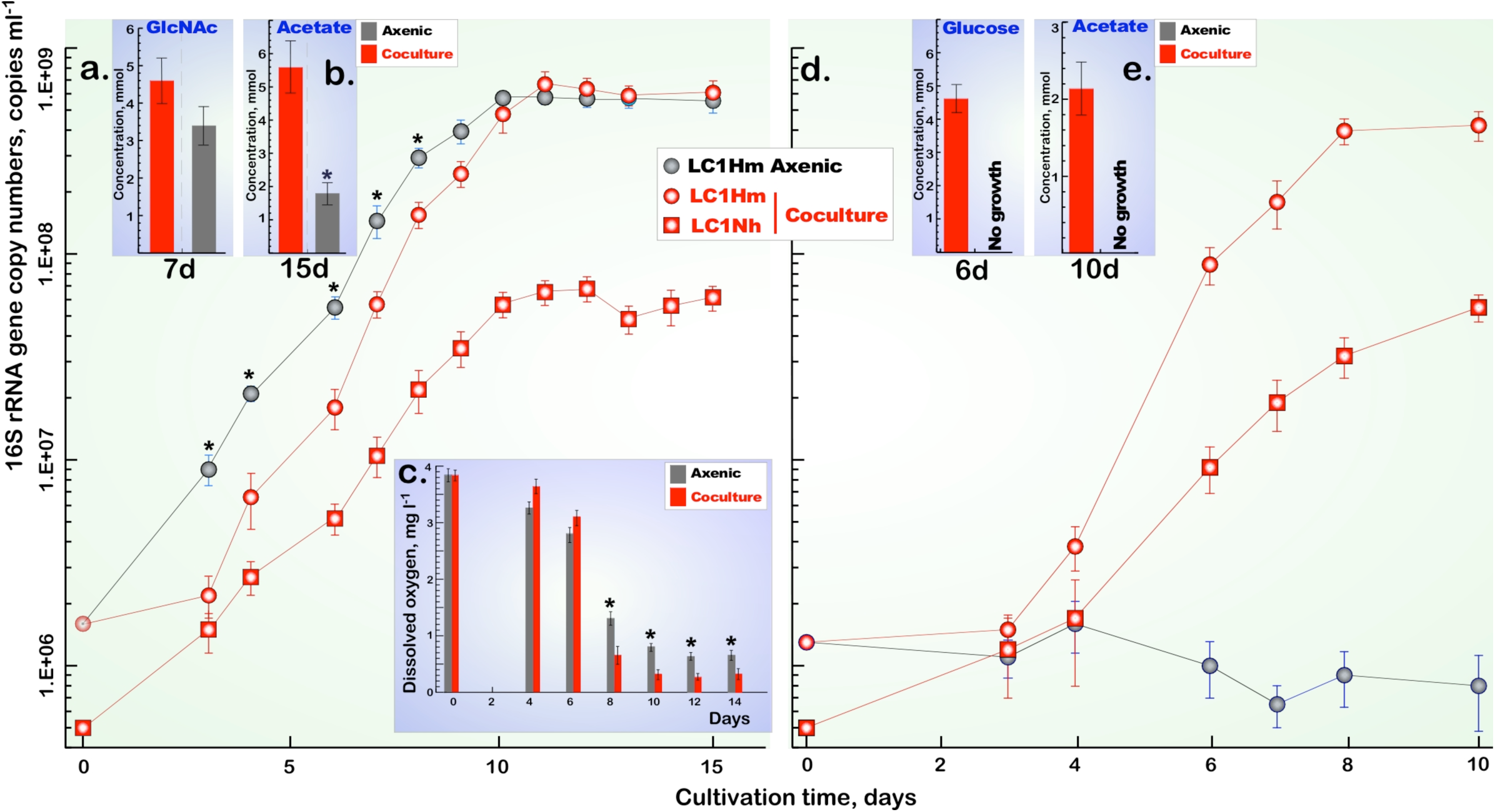
Growth of *Halomicrobium* sp. LC1Hm in pure (axenic) culture and in co-culture with *Ca*. Nanohalobium constans LC1Nh. Panel **a**: growth on chitin. Panel **d**: growth on starch. Insertions (**b, e**) indicate the production of acetate and reduced sugars, N-acetylglucosamine and glucose during growth on chitin and starch, respectively. Insert (**c**) indicates the oxygen consumption of axenic LC1Hm and LC1Hm+LC1Nh cultures during growth on chitin. The overall significance level *p* < 0.01 is shown by asterisks. Error bars (standard deviations) are based on three culture replicates.

None of other the polysaccharides tested (cellulose, glycogen, starch and xylan) supported the growth of *Halomicrobium* sp. LC1Hm in pure culture, so it came as a surprise that the host-ectosymbiont co-culture grew rapidly on alpha-glucans, starch and glycogen. The growth rate (µ_max_ = 0.043-0.066 h^-1^) even exceeded those of the same co-culture and the pure *Halomicrobium* culture on chitin (Fig. 4d; Supplementary Fig. 7). As in the case of chitin (a beta-glucan), degradation of the alpha-glucans was not accompanied by release of measurable amounts of oligomers, and the only detected hydrolysis product was monomeric glucose (≤ 4.61 ± 0.81 mmol) (Fig. 4e). From these experiments we conclude that interspecies cross-feeding occurs in the nanohaloarchaeon-*Halomicrobium* co-culture: *Ca*. Nanohalobium constans is obligately dependent on *Halomicrobium* during growth in all conditions, while *Halomicrobium* benefits from the nanohaloarchaeon during growth on the alpha-glucans (glycogen and starch). The experimental evidence that a DPANN ectosymbiont can provide a specific benefit to its host is, as far as we know, novel.

We succeeded in complete assembly and closure of the *Halomicrobium* sp. LC1Hm genome with the DNA isolated from the host-symbiont co-culture and from the host pure culture, with identical resulting sequences. The nanohaloarchaeal genome was also assembled in its entirety from the co-culture DNA sample. For both genomes, an overview of their characteristics is provided in SM (Supplementary Text, Supplementary Tables 1-4; Supplementary Fig. 8, Extended Data Table 1). As other members of class *Halobacteria, Halomicrobium* sp. LC1Hm is likely to use the “salt-in” strategy of maintaining the osmotic balance, i.e., the accumulation of molar-range concentrations of potassium within cells. Among various specific adaptations, this accumulation implies strong constraints on intracellular proteins to accrue negative charges at their surfaces, known as “acidic” proteome. Calculating the median isoelectric point (pI) for all proteins predicted in *Ca.* Nanohalobium constans LC1Nh and *Halomicrobium* sp. LC1Hm, we found similar acidic values, pI 4.32 and pI 4.7, respectively (Supplementary Table 4). Nanohaloarchaeon differs from the host haloarchaeon by reduced frequencies of histidine and proline and preference for serine over threonine and glutamate over aspartate, which also was noticed by Narasingarao *et al*. (2012)^17^ for *Ca*. Nanosalina and *Ca*. Nanosalinarum spp. and appears to be a property shared by all nanohaloarchaea known so far, thus discriminating them from haloarchaea.

The genome of *Ca*. Nanohalobium consists of a single circular chromosome of 973,463 bp with 43.2% of GC molar content and harbours single copies of 5S, 16S, and 23S rRNA genes located in three different loci, as well as 39 tRNA genes. Of the 1,162 annotated protein-coding genes, 392 (33.7%) could be assigned to a to a functional category within the NCBI COG resource and 732 (63.3%) were assigned to the archaea-specific to arCOGs database; more detailed sequence analysis is likely to increase both percentages. Despite substantial genome completeness, indicated by lack of gaps, presence of tRNAs for all amino acids and identification of the complete set of ribosomal proteins, we identified only 177 genes out of 219 that constitute the core set of archaeal orthologous groups^32^. The set of 219 core arCOGs has been derived from the genome analysis of mostly free-living archaea and did not consider any DPANN species. The genomes of nanohaloarchaea, as well as those of other DPANN archaea, are small and must have experienced many gene deletions, so core arCOGs may not be the best approximation of the essential genes in the small archaea. More specifically, the genome analysis of *Ca*. Nanohalobium predicts the lack of the operational genes for *de novo* synthesis of metabolic precursors, such as amino acids, nucleotides, lipids and enzyme cofactors (Supplementary Text), underscoring its reliance on other members of extremely halophilic microbial community. On the other hand, like most other small bacterial and archaeal genomes, including other DPANN members, the nanohaloarchaeon LC1Nh genome has retained complete sets of the core genes involved in chromosome replication, maintenance and transcription, ribosomal proteins and translation factors.

Analysis of the nanohaloarchaeon genome allowed us to identify the genes implicated in the energy flow and key reactions of biomass production. The ectosymbiont lacks genes for xenorhodopsin biosynthesis, which has been found in some nanohaloarchaeal metagenomic assembled genomes (MAGs)^17^. Furthermore, the *Ca*. Nanohalobium genome contained no genes that encode the components of carbon-fixation pathways. All this indicates a heterotrophic lifestyle of this organisms. The absence of membrane-bound proton-translocating pyrophosphatases suggests that *Ca*. Nanohalobium may rely on the A-type ATPase, the Kef-type potassium-hydrogen antiporter and possibly also other as-yet-unidentified systems, functioning as the outward proton-translocating pumps for maintaining chemiosmotic membrane potential (all information on gene IDs is in Supplementary Text and Extended Data Table 1). Considering the cultivation data and also the absence of genes coding for any known components of the tricarboxylic acid cycle and respiratory complexes, such as NADH dehydrogenase, functional cytochrome oxidases and terminal reductases, *Ca*. Nanohalobium can be classified as an aerotolerant sugar-fermenting anaerobe (Fig. 5; Supplementary Text; Supplementary Table 5).

**Figure 5.**
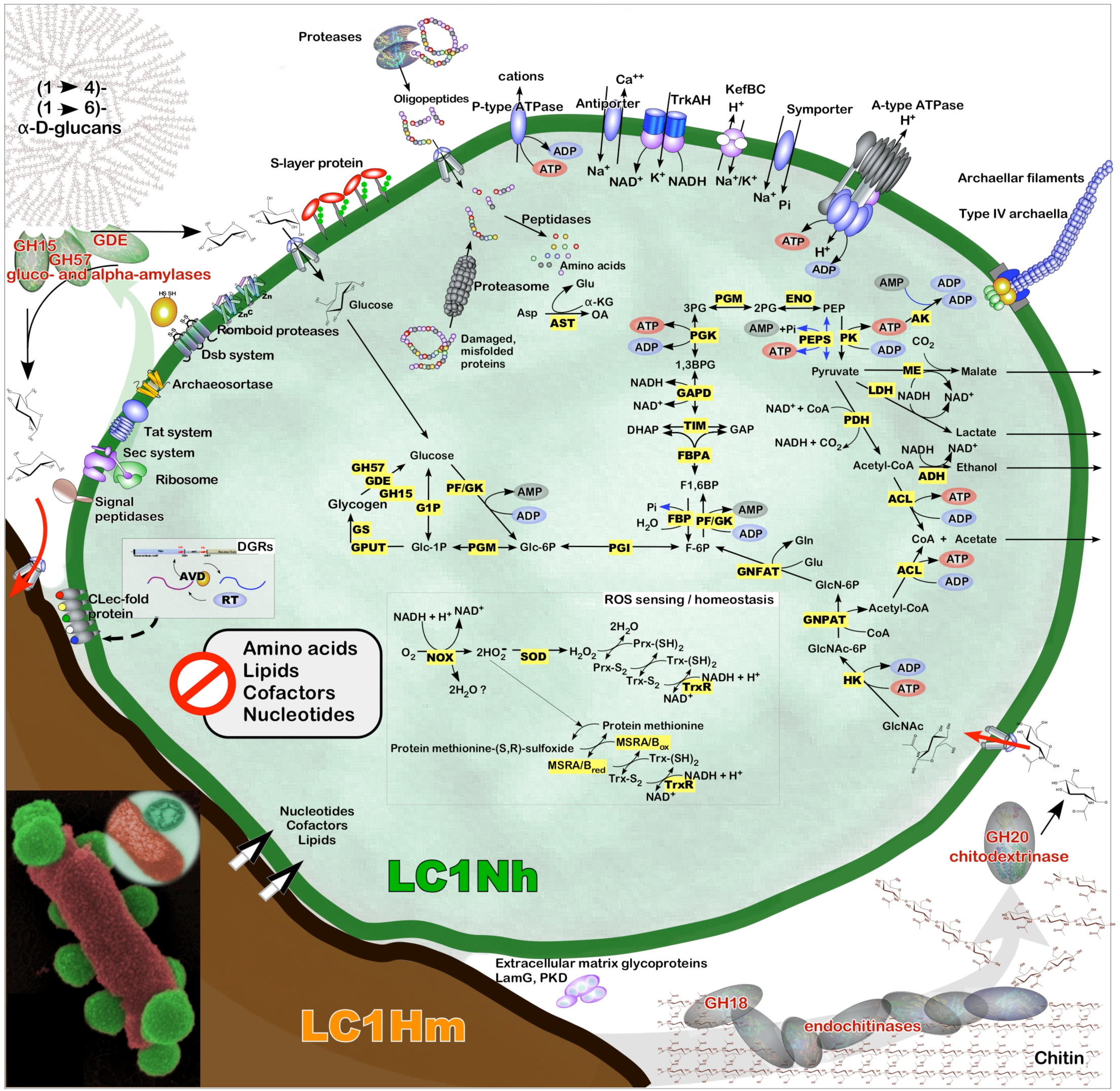
Reconstruction of central metabolic and homeostatic functions of ‘*Ca*. Nanohalobium constans’ LC1Nh based on genomic, proteomic, targeted metabolomics and physiological analyses. Enzymes involved in energy production and in reactive oxygen species (ROS) homeostasis/redox regulation are highlighted in yellow. Chitin degradation enabled by seven extracellularly expressed GH18 endochitinases and one GH20 chitodextrinase of the host, *Halomicrobium* sp. LC1Hm, is shown by grey arrow in the bottom-right part of the figure. Depolymerization of (1→4)- and (1→6)-α-D-glucans by two experimentally confirmed extracellularly expressed glucoamylases of ‘*Ca*. Nanohalobium constans’ LC1Nh is shown with green arrow in the upper-left part of the figure. The “mutualistic” uptake of formed sugars is shown by red arrow. Details on genes and systems abbreviations are provided in the Supplementary Table 5. CoA, coenzyme A; Glc-6P, glucose-6-phosphate; Glc-1P, glucose-1-phosphate; GlcNAc, N-acetyl-glucosamine; GlcNAc-6P, N-acetyl-glucosamine-6-phosphate; GlcN-6P, N-glucosamine-6-phosphate; Glu, glutamate; Gln, glutamine; F-6P, fructose-6-phosphate; F1,6BP, fructose-1,6-biphosphate; DHAP, dihydroxyacetone phosphate; GAP, glyceraldehyde-3-phosphate; 1,3BPG, 1,3-biphosphoglycerate; 3PG, 3-phosphoglycerate; 2PG, 2-phosphoglycerate; PEP, phosphoenol pyruvate; Prx, peroxiredoxin; Trx, thioredoxin.

The chitinolytic activity of the host appears essential for providing energy needs of the ectosymbiont. In agreement with the cultivation results, and as confirmed by omic data, the main path of energy production in nanohaloarchaeon begins with uptake of exogenous GlcNAc formed by chitin-degrading host. Chitinolysis by the host must be mediated by an extensive enzymatic apparatus encoded by multiple loci comprising seven glycosyl hydrolases (GHs) from class III of GH18 family of endochitinases (EC 3.2.1.14) (Supplementary Table 6). Six of them possess N-terminal Tat secretion signals and ChtBD3 chitin-binding domains but no transmembrane helices, indicating that they likely to be extracellular, suitably positioned for digestion of environmental chitin. The simultaneous expression of these extracellular endochitinases (confirmed by the proteomic identification among secreted proteins) indicates that they may have distinct biochemical properties and act synergistically to break down the chitin microfibrils at internal sites. Terminal degradation of chito-oligosaccharides to GlcNAc is expected to occur by the action of extracellular exochitodextrinase, annotated as β-*N*-acetylglucosaminidase of GH20 family (EC 3.2.1.52), which contains the N-terminal Tat signal peptide, a ChtBD3 chitin-binding domain and PKD repeat. Access to high amounts of exogenous GlcNAc appears to be essential for the nanohaloarchaeon, as it has a full set of enzymes responsible for importing and transforming this monosaccharide into fructose-6-P that can then enter central carbohydrate metabolism. All enzymes of glycolysis, gluconeogenesis and glycogen synthesis/catabolism that are encoded by the nanohaloarchaeal genome were detected in cellular proteome and secretome of co-culture when growing with chitin as the carbon source (Supplementary Text, Supplementary Table 5; Extended Data Table 2). Besides the glycogen-debranching enzyme, the genome of nanohaloarchaeal ectosymbiont encodes alpha-amylase and two glucoamylases. The extracellular location of these proteins likely enables the association to utilize exogenous alpha-glucans starch and glycogen as carbon sources alternative to chitin. This extracellular hydrolytic capability, never seen before in DPANN superfamily members, offers new perspective on host-ectosymbiont interaction. Firstly, the ability of *Ca*. Nanohalobium to synthesize and metabolize glycogen is advantageous, because consumption of both intracellular and extracellular (from dead congeners) glycogen would provide at least temporary energetic independence for ectosymbionts dissociated from their host. Secondly, under natural conditions, such glucans may also be provided by halophilic green algae in the form of extracellular linear (1→4)-α-D-glucan polysaccharides^33^. Keeping in mind that arthropods of the genus *Artemia* are less tolerant to extreme salinities (>4M NaCl) than some halophilic green algae such as *Dunaliella*, the association between the nanohaloarchaeon and haloarchaea may benefit the host under conditions when arthtropod chitin is less available than algal derived starch. This may be the case, for example, in crystalliser ponds of solar salterns, or in hypersaline lakes that approaching salt saturation state.

On the one hand, there is evidence for extensive trafficking of nutrients, metabolites and precursors of biopolymers from the *Halomicrobium* host to nanohaloarchaeon. On the other hand, these two tightly associated species do not appear to be involved in rampant gene exchange. Unlike acidophilic members of DPANN superphylum, which have several clusters of orthologous genes shared between the ectosymbiont and the host^24,34^, the nanohaloarchaeal genome shows no synteny with the host genome except for the ancestrally collinear loci coding for ribosomal proteins, suggesting no recent transfer of chromosomal fragments between the two (Supplementary Fig. 9). When all 1162 predicted proteins of *Ca*. Nanohalobium LC1Nh was compared with currently available (at 31 July 2019) nanohaloarchaeal and other DPANN predicted proteomes (Supplementary Fig. 9, Supplementary Text), the vast majority of them (99.2%) were most similar to nanohaloarchaeal proteins, and only five ectosymbiont’s hypothetical proteins were found to be more similar to those of *Halomicrobium* sp. LC1Hm. Other evidences suggest that the genome of *Ca*. Nanohalobium is insulated from horizontal gene transfers to a considerable degree. First, applying the same strategy of isolation to brine-sediment suspension of crystallizer pond of geographically remote solar salterns Margherita di Savoia (Bari, Italy), we obtained a second co-culture consisting of a chitinotrophic *Halomicrobium* sp. CAF1 and a nanohaloarchaeon *Ca*. Nanohalobium constans 2CAF (MN178624). We found that the two distinct nanohaloarchaeal isolates possess virtually identical genomes without any gene-scale insertions, in contrast with much more diverged *Halomicrobium* host isolates (manuscript in preparation). Second, we have found no evidence of integrated viral sequences or recognizable mobile elements in the *Ca*. Nanohalobium genome. Third, the ectosymbiont genome lacks CRISPR-like repeats and encodes none of CRISPR-associated proteins, which are widespread in archaea, including a subset of DPANN genomes^35^. All this suggests that viral infections and acquisition of mobile elements were rare in the evolutionary past of the candidate genus Nanohalobium, possibly reducing the need for innate immunity mechanisms.

A recent study of co-cultures of Antarctic nanohaloarchaea (*Ca*. Nanohaloarchaeum antarcticus) with heterotrophic haloarchaeon *Halorubrum lacusprofundi* shows several remarkable parallels with the present work^21^. Like *Ca*. Nanohalobium LC1Nh, the Antarctic nanohaloarchaea require the presence of a specific haloarchaeon for growth and are predicted to synthesize and metabolize glycogen. Specific trophic and genetic interactions within *Ca*. Nanohaloarchaeum antarcticus – *H. lacusprofundi* consortium remain to be dissected, but even so, these observations raise an important question about the evolution of the archaeal lifestyle (Supplementary Fig. 10; Supplementary Text). Apparently, geographically and phylogenetically diverse members of the *Ca*. Nanohaloarchaeota have evolved specific symbiotic interactions with their hosts, but may have recruited distinct systems for recognizing hosts conferring host specificity. To wit, *Ca*. Nanohaloarchaeum antarcticus possesses unusually large SPEARE proteins, allegedly implicated in attachment to host. Homologs of these genes are found in many nanohaloarchaeal genomes but are missing in *Ca*. Nanohalobium constans, which, in turn, harbours the diversity-generating retroelements, not seen in other nanohaloarchaeal genomes (Supplementary Fig.11 and Supplementary Text). On the other hand, a feature common for all nanohaloarchaea with reconstructed genomes and for other representatives of DPANN superphylum^14^ is the presence of specific lectin-like proteins containing one or more of the laminin globular- or concanavalin A-like domains. These large proteins possess many N-glycosylation sites, are predicted to be on cell surface or extracellular and appear to have a variety of roles including cell adhesion^14^; it is possible that this shared system is a common determinant of host cell attachment system in many Nanohaloarchaeota.

## Discussion

Members of the DPANN superphylum are abundant and globally widespread, but only a handful of representatives of the group have been grown in the laboratory, mostly in binary associations or mixed cultures. The studies of the metabolic capacities, nutritional requirements and ecological roles of DPANN archaea are in their infancy. In this work, we have obtained a binary culture of a nanohaloarchaeon, *Ca*. Nanohalobium constans LC1Nh, with its haloarchaeal host, *Halomicrobium* sp. LC1Hm, and inferred the ecological and physiological basis of this interaction. Association with the host was essential for the growth of nanohaloarchaeon, which showed a strict preference for chitinotrophic haloarchael host of the genus *Halomicrobium*. The nature of the association between the ectosymbiont and its host depends on the type of feedstock: with beta-glucan chitin as the main nutritional source, the nanohaloarchaeal symbiont subsists on the host without conferring any obvious advantage to it, but when the diet is changed to alpha-glucans, the association becomes beneficial to the host. Depending on the evaporation state of hypersaline ecosystems, followed by population dynamics of algae and arthropods, the availability of chitin and alpha-glucans may change multiple times throughout the year; under these circumstances, the lineages of *Halomicrobium* that tolerate the ectosymbiont may be favoured over those that protect themselves against it. The affinity of at least some individuals within *Halomicrobium* populations to colonization by the *Ca*. Nanohalobium cells may be seen as a bet-hedging trait^37,38^, which may reduce the fitness of the host under one condition – in this case, the abundance of beta-glucans – but increase the fitness under another condition, i.e., abundance of alpha-glucans, thereby maximizing the long-term fitness (more precisely, geometric mean fitness). Typically of prokaryotic traits, this candidate bet-hedging trait satisfies the criteria of Category IV evidence^37,38^. Experimental tractability of the host-symbiont system described in this paper may provide more rigorous, Category V and Category VI, evidence of bet-hedging in Archaea. Nanohaloarchaea have small genome sizes and are dependendent on other archaea for growth and reproduction. Our study suggests that these diminutive microbes nevertheless play an active role in the ecology of hypersaline habitats, providing the modality responsible for degradation of abundant biopolymers by proxy: via beneficial interaction with extremely halophilic Euryarchaeota.

## Materials and Methods

### Detailed procedure for sampling

Top 5 cm sediment cores and brine samples were collected from *Saline della Laguna* solar saltern system (37°51’48.70”N; 12°29’02.74”E), one of the ancient sites of salt production near Trapani (Western Sicily), known since VII century BC (Supplementary Figure 1). Sampling was done in June 2016 from the final crystallizer pond *Laguna 27* (265 g l^-1^ salinity, pH 7.2). At the time of sampling, the brines were dominated by halite and were of pinkish colour due to the presence of dense biomass of haloarchaea. The bottom layer of brine was ultramicroaerobic (0.11 ± 0.04 mg l^-1^ of dissolved oxygen) while the sediments were anoxic below 2 cm and contained millimolar concentrations of acid-labile sulfides (HS^-^/FeS). Brine samples for determining major ion concentrations were collected in 1000 ml dark polyethylene (DPE) vials and stored at room temperature. Alternatively, 100 ml of the samples were acidified to pH 2.0 with 100 mM of HNO_3_, diluted to salinity 35 g l^-1^ and stored in DPE vials at room temperature prior to chemical analyses as described elsewhere^39^. Major elements in brine were as follows (ICP-AES data, concentration in g l^-1^): 86.65 Na^+^, 8.56 K^+^, 0.04 Ca^2+^, 5.54 Mg^2+^, 138 Cl^−^ and 20.90 SO_4_^2−^.

### Detailed procedure for enrichment and cultivation conditions and co-culture isolation

Guided by the hydrochemical data, the mineral medium ‘*Laguna Chitin*’ (LC) was prepared on the basis of ONR7a medium^40^, modified by theaddition of (final concentration, g l^-1^): 200 NaCl; 30 MgCl_2_; 0.5 NH_4_Cl; 5 HEPES. The pH was adjusted to 7.0 by 1 M KOH. Chitin from shrimp shells (Sigma-Aldrich catalog number C9213) was added at the final concentration 5 g l^-1^ to serve as growth substrate. After sterilization the medium was supplemented with 1 ml l^-1^ of acidic trace metal solution, 1 ml l^-1^ vitamin mix^41^ and 50 mg of yeast extract. Five grams of the surface (0-2 cm) sediments were mixed with 500 ml of collected brine and enrichment cultures were initiated by addition of 500 ml of sterile LC medium. The enrichment was incubated at 40 °C in tightly closed 1200 ml glass serum bottle without shaking. After 3 months of incubation, the enrichment culture has turned ultramicroaerobic at the bottom of the bottle (0.08 ± 0.02 mg l^-1^ of dissolved oxygen) with chitin particles acquiring simultaneous pinkish colour, characteristic of attached haloarchaeal growth. Determination of oxygen concentration and redox potential was carried out following previously established protocols^39^. This primary enrichment was subcultured by adding one ml of inoculum, collected from the bottom, to 100 ml of the LC medium, supplemented with K_2_HPO_4_ (400 µmol, final concentration), amorphous chitin (2 g l^-1^, final concentration) and bacteria-specific antibiotics, vancomycin and streptomycin (100 mg l^-1^, final concentration), followed by incubating in tightly closed 120 ml serum bottle under the same conditions, i.e. at 37 °C without shaking. Amorphous chitin was prepared by dissolving 10g of crystalline polymer in 200 ml of concentrated HCl on ice overnight followed by dilution in large volume of ice-cold distilled water with subsequent neutralization and concentration by low-speed centrifugation. The final suspension was adjusted to 5% (w/v) concentration in seawater and sterilized at 120 °C for 20 min in closed bottles. After 4 weeks of cultivation the red colour of enrichment intensified with simultaneouschitin degradation. One additional passaging (1 ml of inoculum to 100 ml of sterilized LC medium) was done before the taxonomic characterization of the enriched chitinolytic community. Difference during cultivation on chitin in LC1Hm growth rates and oxygen consumptions were calculated with one-way ANOVA using SigmaStat software. Relative importance of each treatment group was investigated by pairwise Multiple Comparisons procedures (Holm-Sidak method).

The phylogenetic analysis of Illumina barcoding-produced 16S rRNA gene sequences of this chitinolytic enrichment revealed the significant increase of the initial nanohaloarchaeal population from 1 to 33% of total archaeal community (Supplementary Figure 2). Using the LC liquid medium, amended with chitin and antibiotics (see above), we applied the dilution- to-extinction technique to obtain a pure culture. As confirmed by taxon-specific PCR, the 10^−8^ dilution was the last positive transfer of nanohaloarchaea, while chitinotrophic host was found growing also at the next, 10^−9^, dilution. This procedure was repeated two times and the diversity of obtained enrichment was analysed by conventional 16S libraries construction with either nanohaloarchaea-specific or with archaea-specific primers (Supplementary Table 7) and Sanger sequencing of 48 clones from each of those libraries (see 16S rRNA diversity analysis). Phylogenetic analysis revealed that after three rounds of dilution-to-extinction transfers, a binary co-culture of a nanohaloarchaeum and a chitinolyitc haloarchaeon was obtained.

### Attempts to obtain a pure monoculture of *Ca*. Nanohalobium

To test the possibility of maintaining *Ca.* Nanohalobium as a pure monoculture, the binary association was filtered three times through a 0.45 µm pore filter (Sartorius Stedim Biotech) and the filtrate was checked by a *Halomicrobium*-specific PCR to verify the presence of nanohaloarchaeal cells only. Thereafter, 1 ml of filtrate was mixed with 20 ml of fresh LC medium, separately supplemented with various substrates (N-acetylglucosamine, dextrose, maltose, cellobiose, sucrose, starch, glycogen, chitin [2 g l^-1^, final concentration of each carbon source]). In case that Ca. Nanohalobium would need some unknown metabolites supplied by its host in the co-culture, 1 ml of filtrate was inoculated into 20 ml of 1 week-grown co-culture spent medium, obtained by sequential filtration through 0.45-, 0.22- and twice through 0.1 µm pore filters (Sartorius Stedim Biotech). Additionally, 1 ml of filtrate was inoculated into 20 ml of fresh LC medium supplemented with cell lysates, obtained from either pure *Halomicrobium* sp. LC1Hm culture or the binary co-culture. The cells were grown in 20 ml culture with chitin and collected by centrifugation at 13,000 × g for 15 min at 4 °C. The pellets were resuspended in MilliQ water and sonicated using a Vibracell Bioblock Scientific 75115 Sonicator (Sonics & Materials) with 3 bursts each of 30 sec at cycle duty of 50%. All these growth experiments were done in triplicates, statically cultivated for three weeks at 37°C, and then examined by CARD-FISH and PCR. To verify whether physical cell-to-cell contact with *Halomicrobium* is obligately required for *Ca.* Nanohalobium to proliferate, two types of experiments were performed. First, the growth of pure culture of *Halomicrobium* was obtained in the upper chamber of Millipore^®^ Stericup^®^ filtration system (Merck) (capacity 250 ml, pore size 0.22µm), whereas the lower chamber was inoculated with 10 ml of filtrate and 10 ml of fresh minimal LC medium. In the next 3 weeks, a vacuum pump was briefly applied everyday to transfer 5-10 ml of *Halomicrobium*-grown spent medium upside down to the lower chamber. Second, Maxi D-Tube™ Dialyzer (Merck) with 14 kDa molecular cut-off was filled with 3 ml of *Ca.* Nanohalobium-containing filtrate, closed and submerged in 200 ml of fresh LC medium, previously inoculated with *Halomicrobium* pure culture, left to grow for 3 weeks at 37°C and then examined by CARD-FISH and PCR.

### Analyses of metabolites

Oligosaccharides were analysed by High Performance Anionic Exchange Chromatography with Pulsed Amperometric Detection (HPAEC-PAD) on a ICS3000 system (Dionex, Thermo Fischer Scientific Inc., Waltham, MA) consisting of an SP gradient pump, an electrochemical detector with a gold working electrode and Ag/AgCl as reference electrode, and an autosampler (model AS-HV). All eluents were degassed by flushing with helium. An anion-exchange Carbo-Pack PA-200 column (4 × 250 mm, Dionex) connected to a CarboPac PA-200 guard column (4 × 50 mm) was used at 30°C. The initial mobile phase was 4 mM NaOH at 0.3 ml min^−1^ for 30 min. Then the column was washed for 20 min at 0.5 ml min^−1^ with a solution containing 100 mM sodium acetate and 100 mM NaOH, and equilibrated with 4 mM NaOH. Acetate concentrations were measured by gas chromatography (Chromoteq-Crystall 5000.2; column Sovpol-5, 1 m, detector PID in the range of temperatures between 180 °C and 230 °C) after cell removal and acidification of the supernatant to pH 4.0. Additionally, short-chain organic acids acetate, lactate and malate were determined in enrichment supernatants by using corresponding colorimetric assay kits (Sigma-Aldrich catalog numbers MAK086, MAK064 and MAK067-1KT) according to manufacturer’s protocols.

### 16S rRNA diversity analysis

Small aliquot (2 ml) of grown chitinotrophic enrichments was centrifuged at 10,000 × g at room temperature for 10 min. Total DNA was extracted using GNOME DNA kit (MP Biomedicals, USA). The extraction was carried out according to manufacturer’s instructions. Quantity and purity of DNA was estimated by NanoDrop® ND-1000 Spectrophotometer (Wilmington, DE, USA). 16S rRNA gene was amplified using a taxon-specific set of primers for *Nanohaloarchaea*^17^. The reaction was carried out in LifeEco PCR Thermal Cycler (BIOER Technology, China) as follows: initial denaturation at 94 °C for 5 min; 35 cycles of 1 min at 94 °C, 1 min at 60 °C, and 2 min at 72 °C; final extension step of 10 min at 72 °C. PCR products were visualized on an 1% agarose gel. Amplicons for cloning were cut out from the gel and purified with the Wizard SV Gel and PCR Clean–up System kit (Promega, Madison, WI, USA). The other Archaea present in the last positive enrichment were identified following the cloning and sequencing of the 16S rRNA gene using the conventional archaeal universal primers A20F-1492R^42,43^. Library construction was performed as described elsewhere^44^ and 48 positive clones from each libraries were further Sanger-sequenced by Eurofins Genomics (Ebersberg, Germany).

The V3-V4 hypervariable regions of SSU rRNA gene were PCR amplified and the amplicons were sequenced on the Illumina MiSeq platform by FISABIO, Valencia, Spain (http://fisabio.san.gva.es/en/servicios). Library preparations were performed as described previously^45^. The 16S rRNA gene V4 variable region was amplified using PCR primers 341F/785R S-D-Bact-0341-b-S-17, 5’-CCTACGGGNGGCWGCAG-3’ ^46^, and S-D-Bact-0785-a-A-21, 5’-GACTACHVGGGTATCTAATCC-3’ ^46,47^. Barcodes and primer sequences were computationally removed, and then sequences shorter than 150bp, as well as sequences with ambiguous base calls and with homopolymer runs longer than 6bp were removed using ad hoc Python code. Pre-processed sequences were analysed with the NGS analysis pipeline of the SILVA rRNA gene database project (SILVAngs 1.3)^48^. Each sequence was aligned using the SILVA Incremental Aligner (SINA v1.2.10) for ARB SVN^49^ against the SILVA SSU rRNA SEED and quality-controlled database^48^.

Reads with less than 50 aligned nucleotides and with more than 2% of ambiguities and/or homopolymers (low quality), and reads with a low alignment quality (less than 50 identity in the alignment, or alignment score less than 40 as reported by SINA) were excluded from the following analysis. After those quality control steps, sequences were dereplicated on a per-sample basis. Identical reads were collapsed and the unique reads were clustered (OTUs 98% similarity) using cd-hit-est (version 3.1.2; http://www.bioinformatics.org/cd-hit)^50^. The reference read of each OTU was classified by a local nucleotide BLAST search against the non-redundant version of the SILVA SSU Ref dataset (release 132, Dec 13, 2017; http://www.arb-silva.de) using blastn (version 2.2.30+; http://blast.ncbi.nlm.nih.gov/Blast.cgi) with default settings^51^. Reads assigned to Nanohaloarchaea were collected. Only OTUs made up by more than 50 reads were included in the phylogeny analysis and were filtered using the Python script filter_fasta.py (http://qiime.org/scripts/filter_fasta.html). For the initial phylogenetic tree, filtered reads and close sequence relatives were aligned using the SILVA alignment tool^26^ and manually inserted in ARB^27^. The sequences were then re-aligned using the latest SILVA databases for ARB (release 132 SSURef NR99). The neighbour-joining algorithm and the Jukes–Cantor distance matrix within the ARB package were used to generate the phylogenetic trees based on distance analysis for 16S rRNA. One thousand bootstrap re-samplings were performed to estimate the robustness of the tree partitions.

### DAPI and CARD-FISH analyses

Co-culture samples were fixed with formaldehyde (FA, 2% v/v final conc., 1 h at room temperature) and cells were collected by gentle filtration onto polycarbonate membranes (25 mm diameter, 0.22 µm pore size, GE Healthcare Bio-Sciences, USA). Filters for counts of reporter deposition coupled with fluorescence in situ hybridization (CARD-FISH) were embedded in 0.1% low-melting agarose (Sigma), dried at 37 °C for 10 min, and dehydrated with 95% ethanol. Cells were further permeabilized with lysozyme (10mg ml^-1^, 1 h) and achromopeptidase (5mg ml^-1^, 30 min) at 37 °C. Intracellular peroxidase was inhibited by treatment with 10 mM HCl at room temperature for 20 min. Following acid treatment, filters were washed with 0.22 µm filtered MilliQ water, dipped in 95% ethanol and air-dried. Filters were cut in sections for hybridization with the oligonucleotide probes. Using previously published protocol for fluorescence in situ hybridization (FISH) analysis applied for Nanohaloarchaea^17^, we did not observe significant hybridization signal. For CARD-FISH analysis, the horseradish peroxidase Nanohaloarchaea-specific probe Narc_1214^17^ was synthesized by Biomers (Ulm, Germany). Detailed information about the probes is reported in Supplementary Table 7. Universal archaeal probe Arc915 was used to specifically quantify haloarchaeal cells, since Nanohaloarchaea possess two mismatches in the probe-specific site (5’-AGGAATTG**<u>A</u>**CGGGGGA**<u>A</u>**CAC-3’). The HRP probes were added at a final concentration of 50 ng DNA µl^-1^. Hybridization conditions were optimized at 46 °C for 3h, as previously described^52^. Amplification was performed at 46 °C for 40 minutes using tyramide Alexa488 and Alexa594 for Narc_1214 and Arc915 respectively. After amplification, filters were washed in 1x PBS, rinsed in Milli-Q water, dehydrated in 96% ethanol and air-dried. Visualization of all cells was done with DAPI mix (4’,6’–diamidino-2-phenylindole, DNA staining dye; final concentration 2 µg ml-1) in a 4:1 ratio of Citifluor (Citifluor Ltd, Leicester, UK) and Vectashield (Linaris GmbH, Wertheim-Bettingen, Germany)^52^. Cell counts were performed on multiple fields per slide, normalizing 16S rRNA-specific probe counts to the total number of cells stained with the DNA-binding dye 4’,6-diamidino-2-phenylindole. At least 200 DAPI cells in a minimum of 10 fields were counted in the AXIOPLAN 2 Imaging microscope (Zeiss, Germany).

### Determination of cell numbers by quantitative PCR (qPCR)

The qPCR method was employed to determine the relative cell densities of *Halomicrobium* sp. LC1Hm and *Ca.* Nanohalobium constans LC1Nh in co-culture grown in hypersaline LC mineral medium supplemented with vitamins, yeast extract, antibiotics and amorphous chitin, at pH 7.0 and 40 °C. The DNA for quantitative PCR was extracted from 2.0 ml of LC1Hm+LC1Nh co-cultures collected at fixed times, corresponding to lag, exponential and early stationary phases of growth using GNOME DNA kit (MP biomedicals, USA). Extracted DNA was dissolved in 50 µl of TE buffer (10 mM Tris-HCl, 1 mM EDTA [pH 7.5]) and quantified using NanoDrop ND-1000 spectrophotometer (Celbio). The quality of the extracted DNA was checked by electrophoresis in a 1.0% agarose gel.

The qPCR was performed with SYBR Green on an ABI Prism 7300 Real-time PCR System (Applied Biosystems). All amplifications were checked for specificity with dsDNA melt curves and samples exhibiting multiple products were not considered in the analysis. Primers based on the sequences of 16SrRNA genes of LC1Hm and LC1Nh were designed using Primer Express software, version 2.0 (Applied Biosystems, Foster City, Calif.) and listed in the Supplementary Table 8. All primers were checked for specificity using BLAST searches for short, nearly exact matches. All of the sequences in GenBank database, except the desired targets, had ≥ 2 mismatches with the primers. Additionally, the primers were screened for specificity *in silico* using ProbeCheck^53^.

To obtain DNA standards for exact quantification, single clones harboring either *Ca.* Nanohalobium constans LC1Nh or *Halomicrobium* sp. LC1Hm 16S rRNA genes in pGEM-T Easy vector (Promega, Madison, WI, USA) were grown overnight at 37 °C and plasmid was subsequently purified using the NucleoBond Xtra Midi kit (Macherey-Nagel). Plasmid DNA was solubilized in 150 µl of TE buffer and quantified using NanoDrop ND-1000 spectrophotometer (Celbio). With the molecular weight of the plasmid and insert known, tenfold serial dilution series of pGEM_Nh and pGEM_Hm, ranging 3.76 × 10^3^ to 3.76 × 10^9^ copies µl^-1^ and 1.07 × 10^3^ to 1.07 × 10^9^, respectively, were prepared. Serial dilutions were prepared at the same time for each target and used for real-time quantification in triplicate to create the standard curve for sample quantification. Primer concentrations were chosen to minimize the length of quantification cycle, or C_q_, of the standard, while also to minimizing primer-dimers and non-target amplification, as assessed though post-amplification dsDNA melt curves. Each 25 µl reaction contained 50 ng of DNA, 12.5 µl of SYBR Green Master Mix (ThermoFisher), 200 nM of each primers. The qPCR protocol included the following steps: an initial denaturation step at 95 °C for 10 min, followed by 45 cycles of denaturation at 95 °C for 15 s and annealing/elongation at 60 °C for 60 s. A dissociation step was added to check for primer-dimer formation. From the slope of each curve, PCR amplification efficiency (E) was calculated according to the following equation^54^: E = [10^−1/slope^] −1. Obtained slope values fell within the optimal range corresponding to an efficiency of 99.6% and 96.2%, respectively. Detected target genes were converted to cell density (cell ml^-1^) assuming that both organisms are monoploid (possessing of only one chromosome copy).

### Genome sequencing and assembly

Whole-genome shotgun sequencing of the LC1 co-culture was done by FISABIO (Valencia, Spain) using the Illumina MiSeq system platform (San Diego, CA, United States) with 2 × 300 bp short insert paired-end libraries (MiSeq reagent Kit v3). FISABIO also performed the quality assessment by using prinseq-lite program^55^, and the sequence joining (forward R1 and reverse R2) with FLASH program^56^. In the former, the following parameters were applied: min_length = 50; trim_qual_right = 30; trim_qual_type = mean; trim_qual_window = 20. In the latter, default parameters were used. The resulting reads (8,479,286 total paired in two runs and after performing quality check) were assembled by Unicycler 0.4.6 program^57^ with default parameters and using the provided joined sequences as long reads (-l command option). Velvet 1.2.10^58^ plugin inside Geneious package, and Geneious 7.1 built-in assembly software (Biomatters Ltd., New Zealand) were used to control and refine contigs generated from Unicycler, yielding about 147× of LC1Nh and 83× of LC1Hm genome coverage respectively. For closing the LC1Hm genome, a pure isolate sample of LC1 host was further sequenced with the MinION Oxoford Nanopore Technologies platform (Oxford, United Kingdom), yielding the necessary long reads (11,197 total, with 10,371 bp of mean and 103,876 bp of maximum length) to perform a hybrid assembly by Unicycler.

### Gene prediction and annotation

Prediction of *Ca.* Nanohalobium constans LC1Nh protein-coding genes was performed by Glimmer 3.02^59^, and verified by the EMBOSS 6.5.7^60^ inside Geneious 7.1. The FgenesB server^61^ (www.softberry.com) was used for operon prediction. Predicted protein sequences were manually annotated by integrating the results of of blastx/blastp searches against NCBI NR database and information from the databases including KEGG, Clusters of Orthologous Groups (COGs), arCOGs^62-64^ (http://www.genome.jp/tools/blast/), as well as PATRIC/RAST server^65^, and BlastKOALA^66^ (https://www.kegg.jp/blastkoala/). For prediction of tRNA and rRNA genes, tRNAscan-SE 2.0 online^67^ and RNAmmer 1.2 online^68^ were used. Carbohydrate-active enzymes, glycoside hydrolases in particular, were detected by using NCBI blastp against the Carbohydrate Active Enzymes database^69^ (CAZy, http://www.cazy.org/). Transmembrane prediction regions were performed by TMHMM program^70^ (http://www.cbs.dtu.dk/services/TMHMM/). N-glycosylation sites were predicted by NetNGlyc 1.0 Server (http://www.cbs.dtu.dk/services/NetNGlyc/). Genomic comparisons were visualized using Circos software^71^. Percentages of amino-acid identity levels used as the input for Circos visualization were obtained by the RAST server.

### Protein-based phylogeny

The list of 122 archaeal core genes was taken from Genome Taxonomy Data Base (GTDB)^29^. These marker genes were identified in selected genomes using Prodigal v2.6.3^72^, concatenated and then aligned using MAFFT v7.427^73^. Alignment was automatically trimmed using trimAl 1.2rev59^74^. Phylogenetic tree was built using PhyML 3.0 program and the Bayesian-like transformation of approximate likelihood-ratio test for branch support^75^. The substitution model for phylogenetic inference was automatically selected by SMS algorithm^76^. Nanohalorachaeal genomes with completeness < 65% were omitted from analysis. The following non nanohaloarchaeal genomes were included: *Nanoarchaeota* (GCA_000008085.1 and GCA_001552015.1); *Parvarchaeota* (GCA_002498685.1, GCA_002502045.1, GCA_002502525.1, GCA_002503215.1, GCA_002503305.1, GCA_002503915.1 and GCA_002503995.1); *Woesearchaeota* (GCA_000830295.1, GCA_002688315.1, GCA_002688925.1, GCA_002762785.1, GCA_002762985.1, GCA_002779235.1, GCA_000806095.1, SRX764834, SRX764704, UBA10171, UBA12494); *Aenigmarchaeota* (GCA_000806115.1, GCA_002688965.1, GCA_002784265.1, GCA_002789635.1, GCA_002791855.1 and UBA10154). Members of phylum *Micrarchaeota* (GCA_000830275.1, GCA_000402355.1, GCA_002778455.1, GCA_002778455.1, GCA_001871475.1 and SRX764827) were used as an out-group. *Concatenated ribosomal protein phylogeny.* For 16 off the 18 nanohaloarchaeal genomes deposited in NCBI database we constructed another phylogeny. The proteins were concatenated in the following order: 30S ribosomal protein S2 (arCOG04245); 30S ribosomal protein S5 (arCOG04087); 30S ribosomal protein S7 (arCOG04254); 30S ribosomal protein S8 (arCOG04091); 30S ribosomal protein S9 (arCOG04243); 30S ribosomal protein S11 (arCOG04240); 50S ribosomal protein L1 (arCOG04289); 50S ribosomal protein L5 (arCOG04092); 50S ribosomal protein L6 (arCOG04090); 50S ribosomal protein L13 (arCOG04242); 50S ribosomal protein L18 (arCOG04088). The resulting amino acids sequences were aligned by Clustal W 2.1 program with BLOSUM substitution matrix^77^. The tree was inferred by PhyML 3.0 inside Geneious 7.1 with Blosum62 substitution model and 1,000 bootstrap replicates^36^. In this analysis, *Halomicrobium mukohataei* DSM 12286 was used as the outgroup.

### Field emission scanning electron microscopy (FESEM)

Samples were fixed with 2% freshly prepared paraformaldehyde directly in growth medium. Fixative was removed by washing twice with growth medium before final fixation with aqueous osmium tetroxide (4 parts growth medium and 1 part 5% aqueous osmium tetroxide) for 30 min at room temperature. The fixed material was washed with growth medium and placed onto poly-L-lysine coated cover slips for 10 min, followed by treatment with 1% glutaraldehyde to cross-link microbes with L-lysine-coating of the cover slips. This step prevents washing away during the dehydration and critical-point drying of the attached microorganisms. Dehydrating was achieved in a graded series of acetone (10, 30, 50, 70, 90, 100%) on ice for 10 min for each step. Samples in the 100% acetone step were allowed to reach room temperature before another change in 100% acetone. Samples were then subjected to critical point drying with liquid CO_2_ (CPD 030, Bal-Tec, Liechtenstein). Dried samples were covered with a gold-palladium film by sputter coating (SCD 500 Bal-Tec, Liechtenstein) before examination in a field emission scanning electron microscope Zeiss Merlin (Carl Zeiss, Oberkochen) using the Everhart Thornley SE-detector and the inlens-SE detector in a 50:50 ratio with an acceleration voltage of 5 kV. Contrast and brightness were adjusted with Adobe Photoshop CS5.

### Ultrathin sections and transmission electron microscopy (TEM)

Glutaraldehyde-fixed samples (see FESEM) were washed twice with growth medium and fixed with 1% aqueous osmium (final concentration) for 1 hour at room temperature. Samples were then embedded into 2% water agar, solidified agar was cut into small cubes and dehydrated with a graded series of ethanol (10%, 30%, 50%, 70%, 90%, and 100%) for 30 minutes at each step. The 100% ethanol step was repeated twice before the samples were infiltrated with 1 part 100% ethanol and 1 part LRWhite resin for 8 h followed by 1 part 100% ethanol and 2 parts LRWhite resin for 24 h hours, and subsequently infiltrated with pure LRWhite resin with two changes over 2 days. The next day 1 µl starter was added to 10 ml LRWhite resin, stirred and resin was put into 0.5 ml gelatin capsules. The samples were placed into the tip of the capsules, followed by polymerization for 2 days at 50 °C. Ultrathin sections were cut with a diamond knife. Sections were counter-stained with 4% aqueous uranyl acetate for 3 min. Samples were examined in a TEM910 transmission electron microscope (Carl Zeiss, Oberkochen) at an acceleration voltage of 80 kV. Images were taken at calibrated magnifications using a line repliCa. Images were recorded digitally with a Slow-Scan CCD-Camera (ProScan, 1024×1024, Scheuring, Germany) with ITEM-Software (Olympus Soft Imaging Solutions, Münster, Germany). Contrast and brightness were adjusted with Adobe Photoshop CS5.

### Negative staining of whole cells for TEM

Thin carbon support films were prepared by sublimation of a carbon thread onto a freshly cleaved mica surface. Osmium-fixed cells were adsorbed onto the carbon film and negatively stained with 0.5% (w/v) uranyl acetate solution, pH 5.0, according to the method of Valentine et al. (1968)^78^. Samples were examined in a TEM 910 transmission electron microscope (see above).

### Protein digestion

After the centrifugation at 13,000 × g for 15 min at 4 °C of 10 ml chitin-grown LC1Nh+LC1Hm co-culture, cell pellet was dissolved with chaotropic lysis buffer containing 7 M urea (USB Corporation, Cleveland, OH), 2 M thiourea (Sigma-Aldrich), 5 % CHAPS (Sigma-Aldrich), 5 mM TCEP (Sigma-Aldrich) and a protease inhibitor cocktail (Sigma-Aldrich), and incubated for 15 min on ice. Homogenization of the pellet was achieved by ultrasonication for 5 min on ultrasonic bath Branson 2510 (Marshall Scientific, New Hampshire, USA). The homogenate was centrifuged at 20,000 × g for 10 min at 4 °C, and the supernatant containing the solubilized proteins was used for further analysis. The cell-free supernatant obtained after first step of centrifugation was concentrated using a Nanosep 10 kDa cut-off filter (Pall Corporation) to obtain the extracellular (secretory) proteins fraction. Then, 20 µg of proteins in each sample were precipitated by methanol/chloroform method and re-suspended in 30 µl of multichaotropic sample solution UTT buffer (7 M Urea, 2M thiourea, 100 mM TEAB [Sigma-Aldrich]) and 10 µl of 20% SDS was added. Each resuspended sample was reduced with 4 µL of 50 mM TCEP, pH 8.0, at 37 °C for 60 min, followed by addition of 2 µl of 200 mM cysteine-blocking reagent MMTS (SCIEX, Foster City, CA) for 10 min at room temperature. Sample was diluted to 400 µL with S-Trap buffer (90% methanol, 100mM TEAB) and digested into the S-Trap micro column (ProtiFi, NY, USA) according to the manufacturer’s instructions. Digestion was initiated by adding 1 µg of Pierce MS-grade trypsin (Thermo-Fisher Scientific Inc.) to each sample in a ratio 1:20 (w/w), and then incubated at 37 °C overnight on a shaker. Sample digestion was evaporated to dryness in a vacuum concentrator.

### Liquid chromatography and mass spectrometry analysis (LC-MS)

Digested sample was desalted using Stage-Tips with Empore 3M C18 disks (Sigma-Aldrich). 1 µg-aliquot of resulting peptides was subjected to the 1D-nano LC ESI-MS/MS analysis (Liquid Chromatography Electrospray Ionization Tandem Mass Spectrometric) using a nano-liquid chromatography system Eksigent Technologies nanoLC Ultra 1D plus (SCIEX, Foster City, CA, USA) coupled to high-speed Triple TOF 5600 mass spectrometer (SCIEX) with a Nanospray III source. Analytical column used was a silica-based reversed phase Acquity UPLC→ M-Class Peptide BEH C18 Column (Waters Corporation, Milford, MA, USA). The trap column was a C18 Acclaim PepMap™ 100 (Thermo-Fisher Scientific Inc.), 100 µm × 2 cm, 5 µm particle diameter, 100 Å pore size, switched on-line with the analytical column. The loading pump delivered a solution of 0.1 % formic acid in water at 2 µl/min. The nano-pump provided a flow-rate of 250 nL/min and was operated under gradient elution conditions. Peptides were separated using a 250 min gradient ranging from 2 % to 90 % mobile phase B (mobile phase A: 2% acetonitrile [Scharlab S.L.], 0.1 % formic acid [Sigma-Aldrich]; mobile phase B: 100 % acetonitrile, 0.1 % formic acid). Injection volume was 5 µl. Data was acquired using an ion spray voltage floating 2300 V, curtain gas 35, interface heater temperature 150, ion source gas 1 25 and declustering potential 150 V. For IDA parameters, 0.25 s MS survey scan in the mass range of 350–1250 Da were followed by 35 MS/MS scans of 100 ms in the mass range of 100–1800 Da. Switching criteria were set to ions greater than mass to charge ratio (m/z) 350 and smaller than m/z 1250 with charge state of 2–5 and an abundance threshold > 90 counts (cps). Former target ions were excluded for 15 s.

### Proteomics data analysis and sequence search

Mass spectrometry data obtained were processed using PeakView v2.2 Software (SCIEX) and exported as .mgf files. The resulting mass spectra were searched against the predicted peptide sequences encoded by ‘*Ca.* Nhb. constans’ LC1Nh and separately the predicted peptide sequences encoded by *Halomicrobium* sp. LC1Hm (9.394 sequences), together with commonly occurring contaminants, using the open-source software X!TandemPipeline version 3.4.3^79^. Search parameters were set as follows: enzyme, trypsin; allowed missed cleavages, 2; methylthiolation (C) as fixed modification and acetyl (Protein N-term), pyrrolidone from E, pyrrolidone from Q and oxidation (M) as variable modifications. Peptide mass tolerance was set to ± 25 ppm for precursors and 0.05 Da for fragment masses. The confidence interval for protein identification was set to ≥ 95% (p<0.05) and only peptides with an individual ion score above the 1% False Discovery Rates (FDR) at spectra level were considered. Only proteins identified with at least two unique peptides were considered correctly identified. PAI value (protein abundance index) and emPAI, (exponentially modified protein abundance index) were calculated as described elsewhere^80^: PAI = N_obsd_/N_obsbl_, were N_obsd_ and N_obsbl_ are the number of observed peptides per protein and the number of observable peptides per protein, respectively^81^.

## Supporting information

Supplementary Informations

## Data availability

Genomes for *Ca*. Nanohalobium constans LC1Nh and *Halomicrobium* sp. LC1Hm are available under Genbank BioProject PRJNA531595, Biosample SAMN11370769, accession number CP040089. The 16S rRNA gene sequence of *Ca*. Nanohalobium constans 2CAF was deposited in the DDBJ/EMBL/GenBank database under accession numbers MN178624.

## Acknowledgements

We are very grateful and sincerely thank Adele Occhipinti and Luigi Ceci for kind permission and assistance with sampling in solar salterns of Bari and Trapani. We also thank Alexander Yakunin for useful advice and discussion and Alexander Merkel for assistance in phylogenetic analysis. This study was partially supported by grants from the Italian Ministry of University and Research under RITMARE Flagship Project (2012–2016) and of “INMARE” Project (Contract H2020-BG-2014-2634486), funded by the European Union’s Horizon 2020 Research Program. P.N.G. and O.V.G. acknowledge the support from the Centre for Environmental Biotechnology (CEB) Project partly funded by the European Regional Development Fund (ERDF) via the Welsh Assembly Government. D.Y.S. was supported by SYAM-Gravitation Program of the Dutch Ministry of Education and Science (grant 24002002) and by the Russian Foundation for Basic Research (RFBR 19-04-00401). A.R.M. is a Program Director at the National Science Foundation (NSF), the agency of the U.S. Government; his work on this project was supported by the NSF Independent Research and Development Program, but the statements and opinions expressed herein are made in the personal capacity and do not constitute the endorsement by NSF or the government of the United States.

## Author contributions

V.L.C., D.Y.S. and M.M.Y conceived the study. V.L.C., F.C., F.S., D.Y.S. and M.M.Y conducted brine and sediment sampling, cultivation and culture-based experiments. E.M., F.S., S.C., M.F., M.C.M., M.A.S. and S.V.T. performed genome, proteome and metabolite analyses. M.R. and E.A. undertook the microscopy. F.C., R.D., V.L.C. and G.L.S. performed qPCR, SSU rRNA gene analysis and DNA/RNA sequencing. M.M.Y., V.L.C. and F.S. performed chemical analysis. V.L.C., E.M., L.G., P.N.G., O.V.G, D.Y.S., J.E.H., A.M. and M.M.Y. conducted data interpretation. M.M.Y., A.M., D.Y.S., P.N.G., O.V.G. and J.E.H. wrote the manuscript with input from all co-authors. All authors have read and approved the manuscript submission.

## Author information

The authors declare no competing financial interests.

