## Supplementary Informations for "Differential polysaccharide utilization is the basis for a nanohaloarchaeon : haloarchaeon symbiosis"

*"Human subtlety will never devise an invention more beautiful, more simple or more direct than does Nature, because in her inventions, nothing is lacking and nothing is superfluous."*
**Leonardo da Vinci (1452–1519)**

**SUPPLEMENTARY INFORMATION**

### Differential polysaccharide utilization is the basis for a nanohaloarchaeon : haloarchaeon symbiosis.

Violetta La Cono et al.

**Contents** **Page**

**Supplementary Text** **2**

**Supplementary Tables 1 to 8** **13**

**Supplementary Figures 1 to 12** **21**

**References** **33**

**Extended Data Tables 1 and 2 (provided as separated files)**

### Supplementary Text

#### Genome characteristics

The genome of *Ca.* Nanohalobium constans LC1Nh consists of a single circular chromosome of 973,463 bp with GC molar content 43.2%. The chromosome harbours single copies of 5S, 16S, and 23S rRNA genes located in three different loci, as well as 39 tRNA genes, 23 of which have an intron. Of the 1,162 protein-coding genes annotated in LC1Nh, only 392 (33.7%) could be assigned to one of the NCBI COG (Galperin et al., 2015) categories, and 732( 63.3%) to an arCOGs (Makarova et al., 2007, 2015) (Supplementary Tables 1, 2). As of July 2019, there were 19 nanohaloarchaeal genomes of different degree of completeness deposited in NCBI and JGI databases but not included into COG and arCOG resources, and comparisons to these genomes were made as an additional step of the analysis.

The genome of the host haloarchaeon *Halomicrobium* sp. LC1Hm (Supplementary Table 1b) consists of a circular chromosome of 3,105,114 bp, with the GC content 65.7%, two divergent rRNA operons (>95% of gene identity), 48 tRNA genes and two CRISP repeat regions. Of the 3,318 protein-coding genes annotated in LC1Hm, 2,972 were assigned to arCOGs (91.1%). Additionally, *Halomicrobium* sp. LC1Hm had a circular megaplasmid of 223,917 bp, with 64.1% GC content, encoding a single rRNA operon (this has been observed also in the type species of this genus, *Halomicrobium mukohataei* DSM 12286, which has similar genome structure and similar arrangement of rRNA operons [Tindall et al., 2009]) and 183 protein-coding genes. The type species is likely also to be a chitinotroph, judging from the presence of multiple genes encoding endochitinases of the GH18 family (Hou et al., 2014; Sorokin et al., 2015) although that has not been proven in growth experiments.

#### Limited anabolic potential of *Candidatus* Nanohalobium constans LC1Nh.

Similar to most of the currently available DPANN genomes (reviewed in Dombrowski et al., 2019), the 973,463 bp genome of *Ca*. Nanohalobium constans is characterized by the absence of genes encoding the enzymes of canonical anabolic pathways necessary to synthesize most metabolic precursors and intermediates, including purines, pyrimidines, amino acids, cofactors and lipids. The LC1Nh genome is also missing pivotal enzymes of the pentose phosphate pathway, making it incapable of metabolising pentose sugars.

Among the genes of purine and pyrimidine metabolism, we identified in the LC1Nh genome only genes encoding kinases involved in the inter-conversion of nucleoside phosphates (LC1Nh_0358 and 0845). The enzymatic suite for *de novo* amino acid biosynthesis in *Ca*. Nanohalobium constans LC1Nh also is severely limited, with the handful of enzymes in this category, such as asparagine synthase (LC1Nh_0884), aspartate aminotransferase (LC1Nh_0076), threonine dehydratase (LC1Nh_0072), chorismate mutase (LC1Nh_0074, LC1Nh_0078), prephenate dehydratase (LC1Nh_0075) and prephenate dehydrogenase (LC1Nh_0077), mostly representing the downstream stages of synthesis or salvage of amino acids. *De novo* synthesis of cofactors is likewisely nearly absent, with only a few genes completing the synthesis or maturation of the most common cofactors: nicotinamide-nucleotide adenylyltransferase (LC1Nh_0141), riboflavin synthase alpha chain (LC1Nh_0414) and lipoate-protein ligase (LC1Nh_1030). Enzymes involved in C1 turnover are: dihydrofolate reductase (LC1Nh_0153) and 4α-hydroxytetrahydrobiopterin dehydratase (LC1Nh_0691). A caveat in this and other reconstructions is that one-third of the proteins encoded in its genome can not be assigned to any functional category and are annotated as hypothetical proteins, raising the question of whether this nanohaloarchaeon might encode some novel enzymes driving canonical or entirely new metabolic pathways. However, the obligatory host-associated lifestyle of *Ca.* Nanohalobium constans LC1Nh, experimentally validated in this study, is consistent with the observed paucity of important anabolic enzymes, and it is more likely than not that, as with other co-cultured DPANN organisms (Huber et al., 2002; Jahn et al., 2008; Golyshina et al., 2017; Krause et al., 2017; Jarett et al., 2018), nanohaloarchaea must acquire multiple essential metabolites from the host.

**Reconstruction of the central metabolism of *Candidatus* Nanohalobium constans LC1Nh. Expanded text related to Figure 5.**

*Protein translocation systems, membrane-associated cleaving proteases and transporters.* Among 1,205 proteins, predicted in LC1Nh genome, 36 proteins were annotated by BlastKOALA (Kanehisa et al., 2016) as putative members of the *Membrane Transport Category*. Two major complete pathways of protein secretion were found in the LC1Nh genome: the general secretion (Sec) and the twin arginine translocation (Tat) systems. The Sec secretory machinery includes the signal recognition particle complex (SRP-Sec) and five different genes belonging to this protein-trafficking system were recognized: SecE

(LC1Nh_0103), SecY (LC1Nh_1093) SecF/D (LC1Nh_1168-9), the SRP receptor FtsY (LC1Nh_0658) and the targeting protein Ffh (LC1Nh_0652). The Tat pathway is another protein transport system that exports folded proteins from LC1Nh cells. The TatA (LC1Nh_0432) and TatC (LC1Nh_0433) proteins were identified in the LC1Nh genome. Signal peptidases of SppA type (LC1Nh_0849) and of archaeal type I (LC1Nh_0018, 0300 and 0308) are the principal intra-membrane peptidases responsible for processing most of exported proteins in *Ca*. N. constans LC1Nh cells. Two rhomboid-family proteases (LC1Nh_0186, 0519) and archaeosortase (LC1Nh_0663) are the intra-membrane serine proteases that cleave other proteins, including S-layer glycoprotein (LC1Nh_0029, 0824, 1061) within their transmembrane domains. A total of fourteen genes were annotated as components of ABC-type transporter systems. Among them are: a gene encoding for substrate-binding protein (LC1Nh_0508) putatively involved in iron complex ABC transporter system; putative peptide ABC transport system of SalY superfamily (LC1Nh_0028, 0030-1) and three uncharacterized ABC-2 type transporter complexes (LC1Nh_0314-6, LC1Nh_0707-10 and LC1Nh_0762-64). Sugars may be imported into the cytoplasm by one of these ABC- type transporters and/or by major facilitator superfamily (MFS) permease LC1Nh_0802. This putative transporter seems to be highly specific to nanohaloarchaea and has no close homologs with annotated function. Unlike all currently recognized MFS permeases, which have 12 transmembrane helices (TMHs) organised in two domains within a single polypeptide chain, all putative permeases identified in the available nanohaloarchaeal genomes possess only 6-10 TMHs. The sodium-dependent phosphate co-transporter (LC1Nh_0626), zinc/iron permeases of ZIP family (LC1Nh_0011, 1043) along with potassium-dependent sodium-calcium exchanger (LC1Nh_0051), $Na^{+}/K^{+}$: proton antiporter of Kef type (LC1Nh_0771-2,1088), NADH-dependent potassium transport system of Trk type (LC1Nh_0791-2), P-type heavy metal (cations)-transporting ATPase (LC1Nh_0696 and 1022), and K+-dependent mechano-sensitive channel (LC1Nh_1038) likely participate in inorganic ions transportation, osmotic homeostasis and heavy metals resistance.

*Cell-surface structures.* The apparent absence of many anabolic genes suggests that nanohaloarchaeote LC1Nh relies on an external source of metabolic precursors – most likely, on its chitinolytic host organism. Given that, the LC1Nh genome could be expected to contain genes encoding cell-surface structures that would enable its interactions with its host. Similar to what has been described in DPANN relatives of *Ca*. Nanohalobium constans, these interactions are likely mediated by extracellular and/or membrane-associated proteins, including archaella, lectins (carbohydrate-binding proteins) and other proteins that may interact directly with the host cell. At least some of such proteins may be expected to evolve rapidly, in order to overcome the resistance to colonization that the host may develop; thus, any mechanism that is able to generate high diversity is specific loci may be of interest as a potential host-symbiont interaction determinant.

The LC1Nh genome has 21 genes encoding for the archaella assembly machinery and filament proteins (Supplementary Table 5; Extended Data Table 1), and 12 of these genes (LC1Nh_0344-55) are organized in one operon with a highly conserved organization, resembling the characteristic structure of euryarchaeal archaellum operons (Albers and Jarrell, 2018). Scanning electron microscopy of *Ca*. Nanohalobium constans revealed the presence of pilli-like structures similar in appearance and size to protein stalks of the archaella, which can unwind to thin filaments (Figs. 2j,k; Supplementary Figs. 1c-e). It still remains to be seen whether these flagellar structures are used for motility of detached *Ca*. Nanohalobium constans cells, or if they perform an alternative role in attachment to the host cells and/or to polysaccharide substrate.

Transmission and scanning electron microscopy revealed the spatial distribution of up to 17 nanohaloarchaeal cells on a single host cell, as well as what appears to be membrane stretching at the point of contact between the two organisms, suggesting strong intercellular interaction (Figs. 2,3; Supplementary Fig. 5). Given this, we looked for genomic signatures of the mechanisms by which these organism-organism interaction might occur. Besides archaella-related proteins, the LC1Nh genome encodes 11 different glycosyl transferases belonging to families 1, 2 and 8, majority of which are expressed judging from the proteomics data (Extended Data Table 2), indicating that nanohaloarchaeon LC1Nh must expend substantial amounts of produced ATP on the biosynthesis of precusors for glycosylation processes and synthesis of polysaccharides and glycoproteins as principal components of extracellular matrix. Moreover, we detected at least eight strain-specific secreted proteins, containing polycystic kidney disease- (LC1Nh_0257, 0417, 0486, 0919) and concanavalin A-like/lectin- (LC1Nh_0399-401, 0423) domains. Proteins that contain such domains serve a variety of purposes, often mediating interactions with carbohydrates or glycosylated proteins, and predicted to be likely involved in surface interactions in DPANN organisms (Castelle et al., 2018).

*Diversity-generating retroelements (DGRs)*. This family of genetic elements is known to modify DNA sequences and to create massive sequence variation in targeted proteins typically involved in surface attachment, defence and regulation (Medhekar & Miller, 2007). Classical DGR mechanism of mutagenic homing deploys the error-prone reverse transcriptase (epRT) to modify the sequence of the copies of a target protein through diversifications of RNA intermediate. Genes encoding the modified protein contain a variable repeat (VR) in close proximity to an invariant template repeat (TR); epRT-induced mutation of TR-RNA adenines and cDNA replacement of VR leads to an extraordinary degree of sequence diversification. DGRs occur widely in genomes of bacteria and their viruses, and seem to be prevalent in several DPANN lineages (as per GTDB) such as *Nanoarchaeota*, including orders *Pacearchaeales* and *Woesearchaeales*, but not in *Nanohaloarchaeota* (Paul et al., 2017; Dombrowski et al. 2019). DGRs might play a role in cell–cell attachment and providing DPANN organisms with a versatile tool of protein diversification that could be used for adaptation to a dynamic, host-dependent existence by conferring host specificity (Handa et al., 2016; Paul et al., 2017).

The DGRs locus was found in the *Ca.* Nanohalobium constans LC1Nh genome and its enzymatic part consists of the accessory variability determinant (Avd, LC1Nh_0125) and the epRT (LC1Nh_0126) (Supplementary Figure 11). Between the Avd and epRT genes the LC1Nh genome contains a 95-bp long TR, similar to the variable regions (VRA and VRB) of two proteins, LC1Nh_0008 and LC1Nh_0123 (80% and 84%, respectively). At the 3'-end of the VRA, VRB and TR regions, the LC1Nh DGRs system composes of three identical 19 bp-long sequences, coined as initiation of mutagenic homing sequences (IMH). Additionally to these core components, two hairpin/cruciform structures downstream of the VRA and VRB were evident in the LC1Nh genome (Supplementary Figure 11). DGRs hairpin structures were seen to increase the efficiency of genetic information transfer from the TR to the variable regions through an RNA intermediate, a process termed as retro-homing (Guo et al., 2011). The DGR variable proteins LC1Nh_0008 and LC1Nh_0123 are very similar one to another and, as predicted by Phyre2 (Kelley et al., 2015, http://www.sbg.bio.ic.ac.uk/phyre2), belong to the formylglycine-generating enzyme (FGE) subclass with a C-type lectin (CLec)-fold. This finding is in concordance with the recent observation of remarkable conservation in archaea of the ligand-binding CLec-fold for accommodation of massive sequence variation created by DGRs (Le Coq & Gosh, 2011; Handa et al., 2016).

***ROS sensing and redox homeostasis.***

As we discuss in the main text, *Ca.* Nanohalobium constans lacks all respiratory complexes and must have a strictly anaerobic fermentative lifestyle. However, mandatory dependence on aerobic host *Halomicrobium* sp. LC1Hm necessitates an extended tolerance of anaerobic LC1Nh to oxygenated environment. As it discussed in the main text, the host-ectosymbiont co-culture consumed twice as much oxygen as the *Halomicrobium* pure culture, producing microaerobic conditions at the early stationary phase of growth. Increased consumption of oxygen may be due to an increase in metabolic needs of the haloarchaeal host, although the activity of non-respiratory oxygen-scavenging defence systems of nanohaloarchaeon could also also contribute to the elimination of extra oxygen. Analysis of the LC1Nh genome revealed the presence of a thioredoxin system, consisting of two FAD-dependent thioredoxin reductases (TrxR, LC1Nh_0509, 0593), one peroxiredoxin (Prx, LC1Nh_0816) and three thioredoxins (Trx, LC1Nh_0147, 0362, 0828). This sophisticated NAD(P)H-dependent redox system for thiol/disulphide cellular homeostasis in *Ca.* Nanohalobium constans cells may be a part of survival strategy under oxygen exposure, essential for adaptation to aerotolerance. Another finding is the presence in the LC1Nh genome of a putative NADH (peroxi)oxidase (LC1Nh_1142). It is unclear whether this enzyme produces $H_2O$ or $H_2O_2$, but its homolog has been shown to participate in oxygen scavenging and the regeneration of NAD in aero-tolerant anaerobic lactic acid bacteria, which lack the respiratory chains (Geueke et al., 2003). We found also superoxide dismutase (SOD, LC1Nh_0512) and two peptide-methionine sulfoxide reductases specific to each of the forms of the substrate (MSRA and MSRB, LC1Nh_0754 and 0961, respectively), potentially serving to counteract the effect of reactive oxygen species (ROS) produced during metabolism. In addition, we found 4 predicted extracellular dithiol-disulfide isomerases/oxidoreductases (LC1Nh_0053, 0067, 0469, 0814) of DsrA/C family of thioredoxin proteins. In both prokaryotic and eukaryotic organisms these proteins typically participate in oxidative protein folding via disulphide bond formation, breakage and isomerization (Ito and Inaba, 2008). To perform the folding of extracellular proteins, the DsbA/C isomerases/oxidoreductases should be kept in different redox states by interaction with specific membrane-integrated DsbB and DsbG redox regulators. In the LC1Nh genome we found the LC1Nh_0604 protein, which possessed 5 trans-membrane domains and was annotated as disulfide bond formation protein of DsbB family.

**Energy production and catabolism.**

The main genomic features of LC1Nh are consistent with the predictions made on the basis of genome analyses of uncultured nanohaloarchaea and other DPANN organisms. Besides significant genome reduction and presence of full set of genes for chromosome maintenance, these include inability of synthesizing most necessary metabolic precursors, including amino acids and lipids, nucleotides and co-factors. LC1Nh nanohaloarchaeon lacks xenorhodopsin genes, found in other nanohaloarchaeal genomes (Narasingarao et al., 2012), and contains no evidence for carbon fixation pathways, pointing at a strict heterotrophic lifestyle. Given that LC1Nh also lacks genes encoding known components of the tricarboxylic acid (TCA) cycle and any of the respiratory complexes (NADH dehydrogenase, functional cytochrome oxidases and terminal reductases), we infer a strictly anaerobic fermentation-based lifestyle. As the membrane-bound proton-translocating pyrophosphatases were also not detected, the maintenance of a chemiosmotic membrane potential (proton motive force) should rely on the A-type ATPase (LC1Nh_0829-37), the Kef-type potassium-hydrogen antiporter (LC1Nh_0771-2, 1088) and possibly other unidentified systems functioning as the outward proton-translocating membrane pumps.

*Glycolysis*. Complete gene set for the archaeal type of dissimilative Embden-Meyerhof-Parnas (EMP) pathway of glucose degradation was identified in the LC1Nh genome. At the same time, it lacks the Entner-Doudoroff pathway and both oxidative and non-oxidative variants of the pentose phosphate pathways, found in some nanohaloarchaeotes (Ghai et al., 2011; Narasingarao et al., 2012; Finstad et al., 2017; Supplementary Fig. 10). In the absence of membrane respiratory complexes, the EMP pathway is the only way of gaining energy by substrate-level phosphorylation. This central pathway of energy production in nanohaloarchaea shows variations of the upper part also found in some methanogens, including halophilic members, but missing in haloarchaea (Gonzalez-Ordenes et al., 2018). Employing ADP as the phosphoryl donor, the phosphorylation of both glucose and fructose 6-phosphate (F-6P) is likely catalysed by only one enzyme, bifunctional ADP-dependent phosphofructokinase/glucokinase LC1Nh_0114 (PF/GK) of the ribokinase family [EC: 2.7.1.146; 2.7.1.147]. Fructose-1,6-biphosphate is further converted via fructose-biphosphate aldolase LC1Nh_0150 to dihydroxyacetone phosphate and glyceraldehyde-3-phosphate (GAP), which enter the lower portion of the EMP pathway and are further transformed by glyceraldehyde-3-

phosphate dehydrogenase and phosphoglycerate kinase (LC1Nh_0135, 0188), ending with phosphoenolpyruvate (PEP). Of all archaea, only few sugar-utilizing haloarchaea and glycogen-forming methanogens utilize this pair in the glycolytic direction for GAP oxidation and ATP generation (Bräsen et al., 2014). The LC1Nh genome harbours three enzymes capable of catalysing the final step of glycolysis, i.e. conversion of PEP to pyruvate: a pyruvate kinase (LC1Nh_0586) and two AMP/Pi-dependent phosphoenolpyruvate synthases (PEPS, LC1Nh_0232, 1145). Phosphorylations of glucose and F-6P by the PF/GK require 2 molecules of ADP and produce 2 molecules of AMP. Assuming the use of the PGK/GAPDH enzyme pair and pyruvate kinase (LC1Nh_0586), 4 ATPs are generated from 4 ADPs. Adenylate kinase (LC1Nh_0358, 0845) regenerates these ADPs from 2 AMPs and 2 ATPs, resulting in the energy gain of 2 ATP molecules, standard for glycolysis. However, the joint action of bifunctional ADP-dependent PF/GK in the upper part and PEPS in the final step of this modified EMP pathway for sugar degradation could be energetically more favourable (Imanaka et al., 2006; Falb et al. 2008), since produced AMP can be directly re-consumed together with $P_i$, to form ATP, thus resulting favourable energy gain of 4 ATP molecules. As we mentioned above, the genome of *Ca.* Nanohalobium constans encodes a complete archaeal type ATPase complex (9 subunits, LC1Nh_0829-37). Thus, besides many important anabolic and homeostatic reactions, the produced energy can be used for maintenance of cytoplasmic pH within a biocompatible range and for providing a membrane potential by actively pumping protons out of the cytoplasm through the $F_0$ rotor of the A-type ATPase, thus resembling the metabolism of strictly fermentative organisms that lack electron transport chains and incapable of respiration.

*Pyruvate metabolism*. As many DPANN organisms, *Ca.* Nanohalobium constans LC1Nh uses the pyruvate dehydrogenase (PDH) complex LC1NH_0054-57 to decarboxylate pyruvate and to form acetyl-CoA. This complex is frequently found in aerobic bacteria and eukarya, but only a few sugar-utilizing hyperthermophilic and halophilic archaea possess it (Siebers & Schönheit, 2005). ADP-forming acetyl-CoA synthetase (LC1Nh_0059) likely terminates the oxidative pathway of pyruvate metabolism with generation of ATP and formation of acetate. Oxidation of glucose to pyruvate involves the reduction of $NAD^+$ to NADH and thus, to avoid stopping glycolysis, the LC1Nh cells may have to re-oxidize the metabolically unused excess of this reduced electron/energy shuttle. There are multiple indications in the LC1Nh genome that pyruvate could be also used in the reductive pathway as an electron acceptor via NADH-dependent reduction by either lactate dehydrogenase (LC1Nh_0514), or NAD-dependent malic enzyme (LC1Nh_0063), or short-chain alcohol dehydrogenase (LC1Nh_0599, 1170).

*Gluconeogenesis and glycogen metabolism*. The *Ca.* Nanohalobium genome encodes a complete set of enzymes for the archaeal type of gluconeogenesis, including the diagnostic for this pathway bifunctional fructose-1,6-bisphosphate aldolase/phosphatase (LC1Nh_0149). The genome also harbours all key enzymes for glycogen synthesis: phosphomannomutase / phosphoglucomutase (LC1Nh_0113), glycogenin-like protein responsible for the initiation of the glycogen chain (LC1Nh_1199), UTP-glucose-1-phosphate uridylyltransferase (LC1Nh_1188) and glycogen synthase (LC1Nh_0117). This indicates that *de novo* formed glucose, besides being used for glycosylation of various membrane constituents and other metabolic needs, can be stored intracellularly in form of glycogen. This type of carbon and energy storage predicted to be a hallmark of DPANN organisms (Castelle et al. 2015, 2018; Dombrowski et al., 2019), but is absent in all known members of extreme halophilic archaea. Only few methanogens, possessing GAPDH/PGK pair for GAP oxidation, can store glucose in the same way (Bräsen et al., 2014). The capability of glycogen synthesis must be advantageous for *Ca.* Nanohalobium constans LC1Nh, because it would provide at least temporary energetic independence to the cells that dissociate from their host. To decompose glycogen into glucose, LC1Nh encodes glycosyl hydrolases (GHs), such as glycogen debranching enzyme (GDE) / amylo-alpha-1,6-glucosidase (LC1Nh_0116), glucan-1,4-alpha-glucosidase / glucoamylase (LC1Nh_0129) and alpha-amylase (LC1Nh_0131). Remarkably, all these GHs were found both in intracellular and extracellular proteomes (Supplementary Table 5; Extended Data Table 2), indicating that LC1Nh cells might also utilize glycogen from the environment. Joint action of these hydrolases outside the cells would ensure the cleavage of both alpha-1,4 and alpha-1,6 glycosidic linkages, present in exogenous glycogen, producing glucose extracellularly. Glucose can be transported inside the cells by dedicated sugar transporters (the LC1Nh genome harbours three ABC-type transporters of unknown function LC1Nh_0314-6, LC1Nh_0707-10 and LC1Nh_0762-64). Stable growth and maintenance of LC1Nh+LC1Hm co-culture on glucose (5 mmol), used as the single carbon and energy source, confirmed the inferred capability of *Ca.* Nanohalobium constans LC1Nh to uptake this monosaccharide.

**Host-ectosymbiont interactions.**

Our ability to grow nanohaloarchaea in the laboratory allowed us to study many aspects of the trophic network between *Ca.* Nanohalobium constans LC1Nh and *Halomicrobium* sp. LC1Hm, which are likely to be similar to the interactions of the two species that occur in nature. As discussed in the main text, the phenotypic hallmark of pure co-culture is its stable proliferation when insoluble chitin is added as a growth substrate. The detailed analysis of the *Halomicrobium* sp. LC1Hm genome will be presented elsewhere, and only its chitinolytic potential is discussed in this study. This haloarchaeon contains high number of CAZYmes genes, including 26 various glycosyl hydrolases (GHs) (Supplementary Table 6). Among them, seven GHs were unambiguously assigned to class III endochitinases of GH18 family (EC3.2.1.14). All of them are predicted by SignalP 4.0 to have the N-terminal secretion signals, and all of them contain the ChtBD3 chitin-binding domains, suggesting that they may be able to attach to extracellular chitin particles and digest them. According to the CAZY classification, this type of endochitinases breaks down chitin microfibrils at internal sites forming low molecular weight chitodextrins/oligosaccharides $(GlcNAc)_{2-6}$. The *Halomicrobium* sp. LC1Hm genome also encodes a GH20 family protein Hmb_0796, annotated as β-*N*-acetylglucosaminidase (EC3.2.1.14), which could hydrolyze chitodextrins and produce *N*-acetyl-β-glucosamine (GlcNAc). Similar to enochitinases, this enzyme may be acting outside the cells, as indicated by the presence of the signal peptide and two chitin-binding ChtBD3 domains at the N-terminus. The completely extracellular hydrolysis of chitin to GlcNAc by *Halomicrobium* sp. LC1Hm was confirmed by chromatographic analysis of its culture supernatant. Concentrations of GlcNAc were in range of 3.7-5.2 mmol in both chitin-grown axenic culture and in the LC1Nh+LC1Hm co-culture. Such a high amount of exogenous GlcNAc produced by *Halomicrobium* sp. LC1Hm is likely to be advantageous for its nanohaloarchaeal consort, since *Ca.* Nanohalobium constans has full set of enzymes which transform this monosaccharide into fructose-6-P that enters central carbohydrate metabolism. GlcNAc may be imported into the ectosymbiont cytoplasm by one of three ABC- type transporters found in its genome and/or by major facilitator superfamily (MFS) permease LC1Nh_0802. Upon import into the cytoplasm, ATP-dependent kinase (LC1Nh_0180, 1157) can phosphorylate GlcNAc producing GlcNAc-6-P. During two consequent transformations catalyzed by glucosamine-phosphate N-acetyltransferase (LC1Nh_1149, 1150) and glucosamine--fructose-6-phosphate aminotransferase (LC1Nh_0637), GlcNAc-6-P is transformed into F-6-P, thus fuelling both glycolysis and gluconeogenesis. Noteworthy, the phosphorylation of GlcNAc has no energetic costs, since consumed ATP can be regenerated from oxidation of acetyl-

CoA by acetyl-CoA synthetase (LC1Nh_0059) leading to formation of acetate.

Another intriguing feature of *Ca.* Nanohalobium constans is the presence in its genome of at least two extracellular serine proteases (LC1Nh-0159, 0909) and seven different cytoplasmic oligopeptidases (LC1Nh_0029, 0032-35, 0065, 0474) that might utilize exogenous peptides to generate amino acids used to supplement amino acid auxotrophy (Fig. 5 [main text], Supplementary Table 5). This proteolytic suite includes also LC1Nh_0035 glutamyl aminopeptidase of M42 family of metallopeptidases. The same superfamily includes other hydrolases, such as cellulases and endo-1,4-beta-glucanases; this is perhaps the reason that this protein has been annotated in *Ca.* Haloredivivus, *Ca.* Nanopetramus, *Ca.* Nanosalina and *Ca.* Nanosalinarum spp. genomes as cellulase (Ghai et al., 2011; Narasingarao et al., 2012). Our cultivation experiments, however, showed that *Ca.* Nanohalobium constans was unable to grow on cellulose, arguing against the cellulolytic activity of this enzyme. However, the presence of the extracellular proteases might explain the ability of *Ca.* Nanohalobium constans to penetrate the cell wall of its haloarchaeal host observed by the thin section electron microscopy.

**Supplementary Table 1a.** General features of *Ca.* Nanohalobium constans LC1Nh genome.

| Feature | Value |
|---|---|
| Chromosome size | 973,463 bp |
| GC content | 43.2% |
| Protein-coding regions (%) | 883,599 bp (90.8%) |
| Total genes | 1,204 |
| tRNA genes | 39 (23 introns) |
| rRNA genes (5S-16S-23S) | 3 (in 3 different operons) |
| Protein-coding genes | 1,162 |
| Proteins assigned to COGs (%) | 392 (33.7%) |
| Proteins assigned to arCOGs (%) | 735 (66.3%) |
| Average gene length | 760.4 bp |
| Max gene length | 4,500 bp |
| ATG initiation codon proteins | 1,014 |
| GTG initiation codon proteins | 111 |
| TTG initiation codon proteins | 37 |

**Supplementary Table 1b.** General features of *Halomicrobium* sp. LC1Hm genome.

| Feature | Value |
|---|---|
| Chromosome size | 3,105,114 bp |
| GC content | 65.7% |
| Protein-coding regions (%) | 2,736,699 bp (88.1%) |
| Total genes | 3,318 |
| tRNA genes | 48 (3 introns) |
| rRNA genes (5S-16S-23S) | 6 |
| CRISPR regions | 2 |
| Protein-coding genes | 3,264 |
| Proteins assigned to COGs (%) | 2,256 (69.1%) |
| Proteins assigned to arCOGs (%) | 2,972 (91.1%) |
| Average gene length | 833.1 bp |
| Max gene length | 6,555 bp |
| ATG initiation codon proteins | 2,530 |
| GTG initiation codon proteins | 665 |
| TTG initiation codon proteins | 69 |
| Plasmid size | 223,917 bp |
| GC content | 64.1% |
| Protein-coding regions (%) | 196,892 bp (87.9%) |
| Total genes | 187 |
| tRNA genes | - |
| rRNA genes (5S-16S-23S) | 3 |
| CRISPR regions | 1 |
| Protein-coding genes | 183 |
| Proteins assigned to COGs (%) | 130 (71.0%) |
| Proteins assigned to arCOGs (%) | 164 (89.6%) |
| Average gene length | 1,029.7 bp |
| Max gene length | 4,470 bp |
| ATG initiation codon proteins | 138 |
| GTG initiation codon proteins | 43 |
| TTG initiation codon proteins | 2 |

**Supplementary Table 2** Genes of *Ca.* Nanohalobium constans LC1Nh assigned to various functional classes in the NCBI COG database (Galperin et al., 2015).

| **Code** | **Value** | **Percentage** | **COG category** | **Function** |
|---|---|---|---|---|
| J | 92 | 7,92% | Translation, ribosomal structure and biogenesis | Informational |
| K | 23 | 1,98% | Transcription | |
| L | 43 | 3,70% | Replication, recombination and repair | |
| O | 25 | 2,15% | Posttranslational modification, protein turnover, chaperones | Cellular |
| D | 5 | 0,43% | Cell cycle control, cell division, chromosome partitioning | |
| M | 25 | 2,15% | Cell wall/membrane/envelope biogenesis | |
| T | 4 | 0,34% | Signal transduction mechanisms | |
| U | 5 | 0,43% | Intracellular trafficking, secretion, and vesicular transport | |
| V | 17 | 1,46% | Defence mechanisms | |
| C | 18 | 1,55% | Energy production and conversion | Metabolic |
| E | 16 | 1,38% | Amino acid transport and metabolism | |
| F | 15 | 1,29% | Nucleotide transport and metabolism | |
| G | 16 | 1,38% | Carbohydrate transport and metabolism | |
| H | 7 | 0,60% | Coenzyme transport and metabolism | |
| I | 3 | 0,26% | Lipid transport and metabolism | |
| P | 13 | 1,12% | Inorganic ion transport and metabolism | |
| R | 32 | 2,76% | General function prediction | Uncharacterized |
| S | 33 | 2,84% | Function unknown (uncharacterized) | |
| - | **770** | **66,26%** | **Not in COGs** | - |

**Supplementary Table 3.** *Candidatus* Nanohaloarchaeota WGS projects currently registered in the NCBI (JGI) Genome Database. Organisms deposited in GTDB Taxonomy are marked with stars. Complete (circular closed) genomes are in bold.

| Organism name | Accession | Scaffolds | Contigs | Length (bp) | % GC | Isolation Source | Note |
|---|---|---|---|---|---|---|---|
| ***Candidatus* Nanohalobium constans LC1Nh** | **CP040089.1** | **1** | **1** | **973,463** | **43.2** | **Italy: Mozia solar salterns, Trapani** | **Stable cultivation** |
| ***Candidatus* Nanopetramus sp. SG9*** | **CP012986.1** | **1** | **1** | **1,118,574** | **46.4** | **Chile: Salar Grande Halite rock** | **Metagenomic data** |
| ***Cand.* Nanohaloarchaeum antarcticus Nha-CHl** | **Ga0310355 (JGI)** | **1** | **1** | **1,093,273** | **40.3** | **Antarctica: Hypersaline Lake Club** | **Metagenomic data** |
| *Cand.* Nanohaloarchaeum antarcticus Nha-R1 | Ga0101775 (JGI) | 3 | 3 | 1,094,064 | 40.3 | Antarctica: Hypersaline Lake Rauer 1 | Metagenomic data |
| *Candidatus* Haloredivivus sp. G17 | AGNT00000000.1 | 448 | 448 | 1,198,604 | 42.0 | Spain: Santa Pola salterns, Alicante | Single-cell isolation |
| *Nanohaloarchaea* archaeon AB578-D14 | AYGT00000000.1 | 962 | 962 | 1,028,544 | 52.5 | Spain: Santa Pola salterns, Alicante | Single-cell isolation |
| *Candidatus* Nanosalinarum sp. J07AB56 | AEIX00000000.1 | 3 | 259 | 1,215,802 | 55.4 | Australia: Hypersaline Lake Tyrrell | Metagenomic data |
| *Candidatus* Nanosalina sp. J07AB43* | AEIY00000000.1 | 7 | 210 | 1,227,157 | 43.6 | Australia: Hypersaline Lake Tyrrell | Metagenomic data |
| *Nanohaloarchaea* archaeon B1-Br10_U2g1* | LKMN00000000.1 | 26 | 26 | 709,831 | 42.6 | Russia: Kulunda steppe Lake Bitter-1 brine | Metagenomic data |
| *Nanohaloarchaea* archaeon B1-Br10_U2g19* | LKMO00000000.1 | 35 | 35 | 662,884 | 41.0 | Russia: Kulunda steppe Lake Bitter-1 brine | Metagenomic data |
| *Nanohaloarchaea* archaeon B1-Br10_U2g21* | LKMP00000000.1 | 24 | 24 | 815,638 | 39.4 | Russia: Kulunda steppe Lake Bitter-1 brine | Metagenomic data |
| *Nanohaloarchaea* archaeon B1-Br10_U2g29* | LKMQ00000000.1 | 35 | 35 | 526,182 | 40.1 | Russia: Kulunda steppe Lake Bitter-1 brine | Metagenomic data |
| *Nanohaloarchaea* archaeon PL-Br10_U2g16* | LKMV00000000.1 | 54 | 54 | 652,532 | 42.2 | Russia: Kulunda steppe Lake Bitter-1 brine | Metagenomic data |
| *Nanohaloarchaea* archaeon PL-Br10_U2g19* | LKMW00000000.1 | 55 | 55 | 758,571 | 42.4 | Russia: Kulunda steppe Lake Bitter-1 brine | Metagenomic data |
| *Nanohaloarchaea* archaeon PL-Br10_U2g27* | LKMX00000000.1 | 52 | 52 | 581,882 | 43.2 | Russia: Kulunda steppe Lake Bitter-1 brine | Metagenomic data |
| *Nanohaloarchaea* archaeon QH_8_44_6 | PXPB00000000.1 | 329 | 345 | 565,289 | 44.1 | Chile: Atacama Desert salt crust | Metagenomic data |
| *Nanohaloarchaea* archaeon SW_10_44_10 | PXPC00000000.1 | 132 | 144 | 397,623 | 43.3 | Chile: Atacama Desert salt crust | Metagenomic data |
| *Nanohaloarchaea* archaeon SW_4_43_9 | PXPD00000000.1 | 51 | 71 | 661,323 | 43.4 | Chile: Atacama Desert salt crust | Metagenomic data |
| *Nanohaloarchaea* archaeon SW_7_43_1 | PXPE00000000.1 | 1 | 1 | 956,550 | 42.6 | Chile: Atacama Desert salt crust | Metagenomic data |
| *Nanohaloarchaea* archaeon SW_7_46_7 | PXPF00000000.1 | 191 | 224 | 572,429 | 45.7 | Chile: Atacama Desert salt crust | Metagenomic data |

**Supplementary Table 4.** Median isoelectric point (pI) and amino acids composition (%) of all annotated/predicted proteins in DPANN organisms with WGS projects registered into NCBI Genome Database. Organisms validated in GTDB Taxonomy are marked with stars. Members of phylum *Ca*. Nanoarchaeota and the haloarchaeal host, *Halomicrobium* sp. LC1Hm, are shown in blue and gray, respectively..

| **Species** | **pI** | **A Ala** | **C Cys** | **D Asp** | **E Glu** | **F Phe** | **G Gly** | **H His** | **I Ile** | **K Lys** | **L Leu** | **M Met** | **N Asn** | **P Pro** | **Q Gln** | **R Arg** | **S Ser** | **T Thr** | **V Val** | **W Trp** | **Y Tyr** |
|---|---|---|---|---|---|---|---|---|---|---|---|---|---|---|---|---|---|---|---|---|---|
| ***Halomicrobium* sp. LC1Hm** | **4.47** | **11.0** | **0.7** | **8.7** | **8.3** | **3.2** | **8.4** | **2.0** | **3.9** | **1.7** | **8.9** | **1.7** | **2.2** | **4.6** | **2.8** | **6.7** | **5.6** | **6.7** | **9.1** | **1.2** | **2.7** |
| ***C.* Nanohalob. constans LC1Nh** | **4.65** | **6.0** | **0.6** | **7.2** | **11.4** | **4.0** | **7.0** | **1.7** | **6.3** | **6.3** | **8.2** | **2.3** | **4.7** | **3.3** | **3.6** | **4.4** | **6.9** | **5.3** | **6.7** | **0.9** | **3.1** |
| *C.* Nanopetramus sp. SG9* | 4.69 | 6.1 | 0.6 | 7.2 | 11.5 | 4.0 | 7.4 | 1.6 | 5.9 | 6.2 | 8.2 | 2.3 | 4.4 | 3.3 | 3.4 | 4.8 | 6.8 | 5.3 | 6.8 | 0.9 | 3.3 |
| *C.* Haloredivivus sp. G17 | 4.84 | 5.8 | 0.7 | 6.6 | 10.8 | 4.1 | 6.7 | 1.6 | 6.5 | 6.3 | 8.7 | 2.6 | 4.4 | 3.3 | 3.4 | 5.3 | 7.3 | 4.9 | 6.4 | 0.9 | 3.2 |
| *C.* Nanosalinarum sp. J07AB56 | 4.67 | 7.6 | 0.8 | 7.4 | 9.7 | 3.5 | 7.9 | 1.8 | 4.3 | 3.7 | 8.8 | 2.1 | 3.3 | 4.0 | 3.6 | 6.7 | 7.6 | 5.6 | 8.0 | 0.9 | 2.7 |
| *C.* Nanosalina sp. J07AB43* | 4.74 | 5.9 | 0.7 | 7.3 | 10.1 | 3.8 | 6.9 | 1.7 | 6.2 | 5.8 | 8.2 | 2.4 | 4.6 | 3.4 | 3.8 | 5.0 | 7.6 | 5.3 | 6.8 | 0.9 | 3.3 |
| *C.* Nanohal. antarcticus Nha-CHI | 4.84 | 5.6 | 0.7 | 7.1 | 9.9 | 3.9 | 6.9 | 1.7 | 6.8 | 6.3 | 8.5 | 2.5 | 5.0 | 3.4 | 3.8 | 4.6 | 7.1 | 5.4 | 6.6 | 0.9 | 3.3 |
| *C.* Nanohal. antarcticus Nha-R1 | 4.75 | 5.7 | 0.6 | 7.3 | 10.1 | 4.0 | 7.1 | 1.7 | 6.9 | 6.4 | 8.4 | 2.2 | 5.0 | 3.3 | 3.7 | 4.5 | 7.1 | 5.2 | 6.6 | 0.9 | 3.3 |
| *N.* archaeon B1-Br10_U2g1* | 4.74 | 5.8 | 0.6 | 7.0 | 12.0 | 3.8 | 6.8 | 1.7 | 6.4 | 6.4 | 8.6 | 2.3 | 4.1 | 3.3 | 3.2 | 4.9 | 6.5 | 5.2 | 7.2 | 0.8 | 3.3 |
| *N.* archaeon B1-Br10_U2g19* | 4.76 | 5.7 | 0.7 | 7.1 | 11.9 | 3.9 | 6.7 | 1.7 | 6.3 | 6.5 | 8.6 | 2.2 | 4.4 | 3.3 | 3.3 | 4.6 | 6.6 | 5.4 | 7.1 | 0.8 | 3.4 |
| *N.* archaeon B1-Br10_U2g21* | 4.70 | 6.1 | 0.7 | 7.2 | 11.7 | 3.9 | 6.9 | 1.7 | 6.6 | 6.0 | 8.5 | 2.3 | 4.6 | 3.3 | 3.3 | 4.5 | 6.7 | 5.0 | 6.7 | 0.9 | 3.4 |
| *N.* archaeon B1-Br10_U2g29* | 4.75 | 6.0 | 0.6 | 7.1 | 11.4 | 4.2 | 6.9 | 1.6 | 6.7 | 6.2 | 8.5 | 2.5 | 4.5 | 3.3 | 3.3 | 4.6 | 6.8 | 4.9 | 6.9 | 0.9 | 3.2 |
| *N.* archaeon PL-Br10_U2g16* | 4.65 | 5.5 | 0.6 | 7.7 | 12.1 | 3.9 | 7.1 | 1.6 | 6.5 | 6.5 | 8.3 | 2.4 | 4.2 | 3.2 | 3.3 | 4.7 | 6.8 | 4.7 | 6.9 | 0.8 | 3.2 |
| *N.* archaeon PL-Br10_U2g19* | 4.79 | 5.6 | 0.7 | 7.0 | 12.0 | 3.9 | 6.7 | 1.7 | 6.4 | 6.5 | 8.5 | 2.3 | 4.2 | 3.5 | 3.2 | 4.9 | 6.5 | 5.3 | 7.0 | 0.8 | 3.4 |
| *N.* archaeon PL-Br10_U2g27* | 4.90 | 6.4 | 0.8 | 6.7 | 11.3 | 4.0 | 6.9 | 1.7 | 6.4 | 6.4 | 8.9 | 2.4 | 4.1 | 3.4 | 3.3 | 4.8 | 6.7 | 4.9 | 6.9 | 0.9 | 3.2 |
| *N.* archaeon QH_8_44_6 | 4.90 | 5.5 | 0.9 | 6.1 | 10.5 | 3.8 | 6.7 | 1.8 | 6.0 | 6.7 | 8.2 | 2.3 | 4.3 | 3.4 | 3.7 | 6.4 | 7.1 | 5.3 | 6.1 | 1.0 | 3.1 |
| *N.* archaeon SW_10_44_10 | 4.88 | 5.6 | 0.8 | 6.3 | 10.8 | 3.7 | 6.8 | 1.7 | 6.4 | 7.0 | 8.1 | 2.5 | 4.4 | 3.3 | 3.5 | 5.8 | 6.9 | 5.1 | 6.4 | 0.9 | 3.1 |
| *N.* archaeon SW_4_43_9 | 4.81 | 5.7 | 0.8 | 6.6 | 11.2 | 3.8 | 7.0 | 1.6 | 6.5 | 6.8 | 8.2 | 2.3 | 4.6 | 3.3 | 3.4 | 5.3 | 6.9 | 5.2 | 6.5 | 0.9 | 3.2 |
| *N.* archaeon SW_7_43_1 | 4.79 | 5.7 | 0.7 | 6.9 | 11.3 | 3.9 | 7.0 | 1.6 | 6.6 | 6.7 | 8.2 | 2.3 | 4.9 | 3.2 | 3.3 | 4.8 | 6.9 | 5.2 | 6.6 | 0.9 | 3.3 |
| *N.* archaeon SW_7_46_7 | 4.81 | 5.8 | 0.8 | 6.3 | 10.5 | 3.7 | 7.1 | 1.8 | 5.7 | 6.3 | 8.1 | 2.3 | 4.2 | 3.5 | 3.9 | 6.3 | 7.1 | 5.4 | 6.3 | 0.9 | 3.0 |
| *C.* N. archaeon B72_G9 | 8.27 | 6.1 | 1.6 | 5.9 | 6.6 | 4.0 | 6.4 | 2.0 | 8.5 | 8.6 | 8.5 | 2.6 | 4.8 | 3.5 | 2.7 | 4.5 | 6.3 | 5.8 | 6.6 | 0.9 | 3.8 |
| *C.* N. archaeon B49_G9 | 8.84 | 6.0 | 1.6 | 5.2 | 7.0 | 4.7 | 5.9 | 1.7 | 9.7 | 10.2 | 8.4 | 2.6 | 5.7 | 3.2 | 2.6 | 3.8 | 6.2 | 5.0 | 5.5 | 0.8 | 3.9 |
| *Nanoarchaeum equitans* Kin4-M* | 8.81 | 5.2 | 0.8 | 5.0 | 7.9 | 4.4 | 5.3 | 1.3 | 10.5 | 10.8 | 10.4 | 1.7 | 5.3 | 4.0 | 2.2 | 3.9 | 4.6 | 4.1 | 5.9 | 1.0 | 5.6 |
| *C.* Aenigmarchaeota ex4484_224* | 9.30 | 4.5 | 0.9 | 3.9 | 8.8 | 5.7 | 5.5 | 1.2 | 10.7 | 12.2 | 10.1 | 1.8 | 4.6 | 3.4 | 2.1 | 4.5 | 5.8 | 3.5 | 6.0 | 1.1 | 3.8 |

**Supplementary Table 5.** Proteins of the central metabolic and homeostatic reconstruction map described in Figure 5 (see main text).

| Category | locus_tag | Product | Name in Figure 5 | Proteome biomass | Proteome secretome |
|---|---|---|---|---|---|
| **Proteases and peptitades** | LC1Nh_0018,0300 | signal peptidase I | | YES | NO |
| | LC1Nh_0308 | signal peptidase I | | NO | NO |
| | LC1Nh_0032,0033,0035,0065,0159,0474 | peptidase | | YES | YES |
| | LC1Nh_0909 | serine protease Do | | YES | YES |
| | LC1Nh_0076 | aspartate aminotransferase | | YES | NO |
| **Poly- and oligo- saccharides** | LC1Nh_0116 | glycogen debranching enzyme (alpha-1,6-glucosidase) | GDE | YES | YES |
| | LC1Nh_0129 | glycosyl hydrolase family 15 | GH15 | YES | YES |
| | LC1Nh_0131 | glycosyl hydrolase family 57 | GH57 | YES | YES |
| | LC1Nh_0795 | glycosyl hydrolase family 1 | GH1 | YES | YES |
| | LC1Nh_0796 | glycosyltransferase family 2 protein | | YES | YES |
| **NAGA** | LC1Nh_0180,1157 | Gluco(hexo)kinase | HK | YES | YES |
| | LC1Nh_0637 | glucosamine--fructose-6-phosphate aminotransferase | GNFAT | YES | YES |
| | LC1Nh_1149 | bifunctional UDP-N-acetylglucosamine pyroph. | GNPAT | NO | YES |
| | LC1Nh_1150 | bifunctional UDP-N-acetylglucosamine pyroph. | GNPAT | YES | YES |
| | LC1Nh_0802 | major facilitator superfamily MFS_1 | MFS | YES | NO |
| **Glycolysis** | LC1Nh_0114 | ADP-dependent phosphofructokinase/glucokinase | PF/GK | NO | YES |
| | LC1Nh_0115 | triosephosphate isomerase (TIM) | TIM | YES | YES |
| | LC1Nh_0118 | glucose-6-phosphate isomerase | PGI | YES | YES |
| | LC1Nh_0135 | glyceraldehyde-3-phosphate dehydrogenase (NAD(P)) | GAPD | YES | YES |
| | LC1Nh_0150 | fructose-1,6-bisphosphatase I | FBPA | YES | YES |
| | LC1Nh_0188 | phosphoglycerate kinase | PGK | YES | YES |
| | LC1Nh_0232 | phosphoenolpyruvate synthase / pyruvate, water | PEPS | YES | YES |
| | LC1Nh_0282 | enolase, C-terminal TIM barrel domain | ENO | YES | NO |
| | LC1Nh_0358,0845 | adenylate kinase | AK | YES | NO |
| | LC1Nh_0586 | pyruvate kinase | PK | YES | YES |
| | LC1Nh_0954 | 2,3-bisphosphoglycerate-independent phosphoglycer. | PGM | YES | NO |
| | LC1Nh_1033 | glucose-1-phosphatase | | YES | NO |
| **Pyruvate Metab. & Fermentation** | LC1Nh_0054,0055,0056,0057 | pyruvate dehydrogenase (dihydrolipoamide dehydr.) | PDH | YES | YES |
| | LC1Nh_0059 | acetate-CoA ligase (ADP-forming) subunit alpha | ACL | YES | NO |
| | LC1Nh_0063 | malate dehydrogenase (oxaloacetate-decarboxylating) | ME | YES | YES |
| | LC1Nh_0514 | D-lactate dehydrogenase | LDH | YES | NO |
| | LC1Nh_0599,1170 | SDR family oxidoreductase | ADH | YES | YES |
| **Gluconeogenesis** | LC1Nh_1033 | glucose-1-phosphatase | G1P | YES | NO |
| | LC1Nh_0149 | fructose-bisphosphate aldolase, class I | FBP | YES | YES |
| **Glycogen synthesis** | LC1Nh_0113 | phosphomannomutase / phosphoglucomutase | PGM | YES | YES |
| | LC1Nh_0117 | glycogen synthase | GS | YES | YES |
| | LC1Nh_1188 | nucleotidyl transferase | GPUT | YES | YES |
| | LC1Nh_1199 | glycogenin-like protein | GLP | YES | NO |
| **Secretion** | LC1Nh_1093,1168 | preprotein translocase subunit SecY | Sec | NO | YES |
| | LC1Nh_1169 | preprotein translocase subunit SecD | system | YES | YES |
| | LC1Nh_0432 | sec-independent protein translocase protein TatA | Tat | YES | NO |

| | | | | | |
|---|---|---|---|---|---|
| | LC1Nh_0433 | sec-independent protein translocase protein TatC | system | NO | NO |
| **Oxidative stress** | LC1Nh_0147 | thioredoxin family protein | Trx | NO | NO |
| | LC1Nh_0362,0593 | thioredoxin | TrxR | YES | NO |
| | LC1Nh_0509 | thioredoxin reductase (NADPH) | TrxR | NO | NO |
| | LC1Nh_0512 | superoxide dismutase, Fe-Mn family | SOD | YES | YES |
| | LC1Nh_0754 | peptide-methionine (S)-S-oxide reductase | MSRA | NO | NO |
| | LC1Nh_0816 | OsmC family peroxiredoxin | Prx | NO | NO |
| | LC1Nh_0828 | glutaredoxin | | NO | NO |
| | LC1Nh_0961 | peptide-methionine (R)-S-oxide reductase | MSRB | YES | NO |
| | LC1Nh_1142 | pyridine nucleotide-disulphide oxidoreductase | NOX | YES | NO |
| **Cell surface & Flagella** | LC1Nh_0029,0824,1085 | S-layer domain protein | | YES | YES |
| | LC1Nh_0305,0500,0738,0929 | PilT protein domain protein | | YES | NO |
| | LC1Nh_0350,1130 | archaeal flagellar protein FlaI | | YES | YES |
| | LC1Nh_0351,0352,1131 | archaeal flagellar protein FlaJ | | YES | YES |
| | LC1Nh_0941 | Type IV secretory pathway ATPase VirB11/Archaellum | | YES | YES |
| | LC1Nh_1129 | hypothetical protein | | YES | YES |
| | LC1Nh_1132 | archaeal flagellar protein FlaJ | | NO | YES |
| | LC1Nh_0399,0400,0401,0423 | concanavalin A-like lectin/glucanases superfamily | LamG | YES | YES |
| | LC1Nh_0791,0792 | Trk-type K+ transport system, membrane component | TrkAH | YES | YES |
| | LC1Nh_1088 | Kef-type K+ transport system, membrane component | KefBC | YES | YES |
| | LC1Nh_0626 | Na+/phosphate symporter | Symportr | YES | NO |
| | LC1Nh_0008 | formylglycine-generating enzyme | CLec-fold protein | YES | YES |
| **DGR** | LC1Nh_0123 | hypothetical protein | | YES | NO |
| | LC1Nh_0124 | hypothetical protein | | NO | NO |
| | LC1Nh_0125 | diversity-generating retroelement protein bAvd | Avd | NO | NO |
| | LC1Nh_0126 | RNA-dependent DNA polymerase (Reverse transcr.) | RT | YES | YES |
| | LC1Nh_0127 | transcriptional regulator like protein | | NO | NO |
| **Energy** | LC1Nh_0829-0837, LC1Nh_0833-0837 | V/A-type H+/Na+-transporting ATPase subunits | A-type ATPase | YES | YES |
| **Protein translocation and** | LC1Nh_0284,0475,0554,0585 | proteasome | Proteaso | YES | YES |
| **Posttranslational modification** | LC1Nh_0849 | protease IV | me | YES | NO |
| | LC1Nh_0186,0519 | rhomboid family intramembrane serine protease | | YES | YES |
| **PKD-containing proteins** | LC1Nh_0257,0486,0919 | PKD domain containing protein | PKD | YES | NO |
| **Protein-disulfide isomerase /** | LC1Nh_0053,0067,0814 | DSBA oxidoreductase | Dsb | YES | YES |
| **oxidoreductase Dsb system** | LC1Nh_0605 | disulfide bond formation protein DsbB | system | NO | NO |

**Supplementary Table 6.** Proteins identified by CAZy and NCBI databases as glycoside hydrolases (GH) in the *Ca.* Nanohalobium constans LC1Nh and the *Halomicrobium* sp. LC1Hm genomes. Endochitinases and exochitodextrinase of *Halomicrobium* sp. LC1Hm are highlighted in grey.

| locus_tag | NCBI Accession | GH family | Activities in family (most common) | Blastp bitscore | Blastp E-value |
|---|---|---|---|---|---|
| LC1Nh_0116 | AOV94584.1 | GH_NC | Glycoside hydrolases not yet assigned to a family | 645 | 0 |
| LC1Nh_0795 | AHB42465.1 | GH1 | β-glucosidase; β-galactosidase; β-mannosidase | 306 | 2,00E-98 |
| LC1Nh_0129 | PIZ00265.1 | GH15 | Glucoamylase; glucodextranase; α,α-trehalase | 347 | 8,00E-115 |
| LC1Nh_0131 | AOV94591.1 | GH57 | α-amylase; α-galactosidase; amylopullulanase | 714 | 0 |
| LC1Hm_0354 | ACV46754.1 | CBM5 | Cellulose-binding domain family V | 1739 | 0 |
| LC1Hm_2080 | ACV48841.1 | CBM5 | Cellulose-binding domain family V | 752 | 0 |
| LC1Hm_0070 | ACV47013.1 | CBM5, GH18 | Cellulose-binding domain family V, chitinase | 734 | 0 |
| LC1Hm_0809 | ACV47620.1 | CBM5, GH20 | Cellulose-binding domain family V, β-hexosaminidase; lacto-N-biosidase | 1405 | 0 |
| LC1Hm_0822 | ACV47608.1 | CBM5, GH18 | Cellulose-binding domain family V, chitinase | 786 | 0 |
| LC1Hm_2270 | ACV49026.1 | CBM5, GH18 | Cellulose-binding domain family V, chitinase | 976 | 0 |
| LC1Hm_2271 | ACV49027.1 | CBM5, GH18 | Cellulose-binding domain family V, chitinase | 677 | 0 |
| LC1Hm_2272 | ACV49028.1 | CBM5, GH18 | Cellulose-binding domain family V, chitinase | 1088 | 0 |
| LC1Hm_2273 | ACV49029.1 | CBM5, GH18 | Cellulose-binding domain family V, chitinase | 867 | 0 |
| LC1Hm_1158 | BAC76692.1 | GH18 | Chitinase; lysozyme; endo-β-N-acetylglucosaminidase | 271 | 1,00E-84 |
| LC1Hm_0425 | AEH39070.1 | GH2 | β-galactosidase; β-mannosidase; β-glucuronidase | 1531 | 0 |
| LC1Hm_1176 | ADB63454.1 | GH2 | β-galactosidase; β-mannosidase; β-glucuronidase | 952 | 0 |
| LC1Hm_2335 | AAV46707.1 | GH2 | β-galactosidase; β-mannosidase; β-glucuronidase | 663 | 0 |
| LC1Hm_4132 | ACV49329.1 | GH2 | β-galactosidase; β-mannosidase; β-glucuronidase | 1904 | 0 |
| LC1Hm_1210 | AGB37354.1 | GH3 | β-glucosidase; xylan 1,4-β-xylosidase; β-glucosylceramidase | 722 | 0 |
| LC1Hm_2524 | ACV49279.1 | GH3 | β-glucosidase; xylan 1,4-β-xylosidase; β-glucosylceramidase | 974 | 0 |
| LC1Hm_2617 | AKU08087.1 | GH3 | β-glucosidase; xylan 1,4-β-xylosidase; β-glucosylceramidase | 982 | 0 |
| LC1Hm_4049 | AEH38218.1 | GH4 | Maltose-6-phosphate glucosidase; α-glucosidase; α-galactosidase | 667 | 0 |
| LC1Hm_0270 | ACV46974.1 | GH9 | Endoglucanase; endo-β-1,3(4)-glucanase / lichenase-laminarinase; β-glucosidase | 1001 | 0 |
| LC1Hm_1562 | ACV48372.1 | GH13_31 | α-amylase; pullulanase; cyclomaltodextrin glucanotransferase | 1049 | 0 |
| LC1Hm_2614 | ACV46202.1 | GH13_31 | α-amylase; pullulanase; cyclomaltodextrin glucanotransferase | 1036 | 0 |
| LC1Hm_2615 | ACV46203.1 | GH13_31 | α-amylase; pullulanase; cyclomaltodextrin glucanotransferase | 1047 | 0 |
| LC1Hm_1157 | BAZ30250.1 | GH29 | α-L-fucosidase; α-1,3/1,4-L-fucosidase | 289 | 7,00E-90 |
| LC1Hm_1173 | BAZ54052.1 | GH29 | α-L-fucosidase; α-1,3/1,4-L-fucosidase | 347 | 3,00E-112 |
| LC1Hm_1261 | ACV48052.1 | GH32 | Invertase; endo-inulinase; β-2,6-fructan 6-levanbiohydrolase | 1100 | 0 |
| LC1Hm_1177 | AGF92912.1 | GH38 | α-mannosidase; mannosyl-oligosaccharide α-1,2-mannosidase | 916 | 0 |
| LC1Hm_1260 | ACV48051.1 | GH68 | Levansucrase; β-fructofuranosidase; inulosucrase | 830 | 0 |
| LC1Hm_4127 | ACV49500.1 | GH88 | d-4,5-unsaturated β-glucuronyl hydrolase | 764 | 0 |

**Supplementary Table 7.** Details on CARD-FISH probes and conditions

| Probes | Sequences (5'-3') | FA (%) | Hybridization temperature (°C) | Washing temperature (°C) |
|---|---|---|---|---|
| Arch915 | GTGCTCCCCCGCCAATTCCT | 20 | 46 | 48 |
| Narc_1214 | CCGCGTGTATCCCAGAGC | 20 | 46 | 48 |

**Supplementary Table 8.** qPCR primers used in the present study.

| Organism | Primer | 5’ → 3’ | Amplification efficiency (E) |
|---|---|---|---|
| *Ca.* N. constans LC1Nh | Nh_1014F | CGTGAGGTGTCCGGTTAAGT | 99.6% |
| | Nh_1130R | GCTCCTTCCCCTGGTTTATC | |
| *Halomicrobium* sp. LC1Hm | Hm_0409 | TTCTCGACCGTAAGGTGGTC | 99.2% |
| | Hm_0527 | CAAGCTACGGACGCTTTAGG | |

#### Supplementary Figures

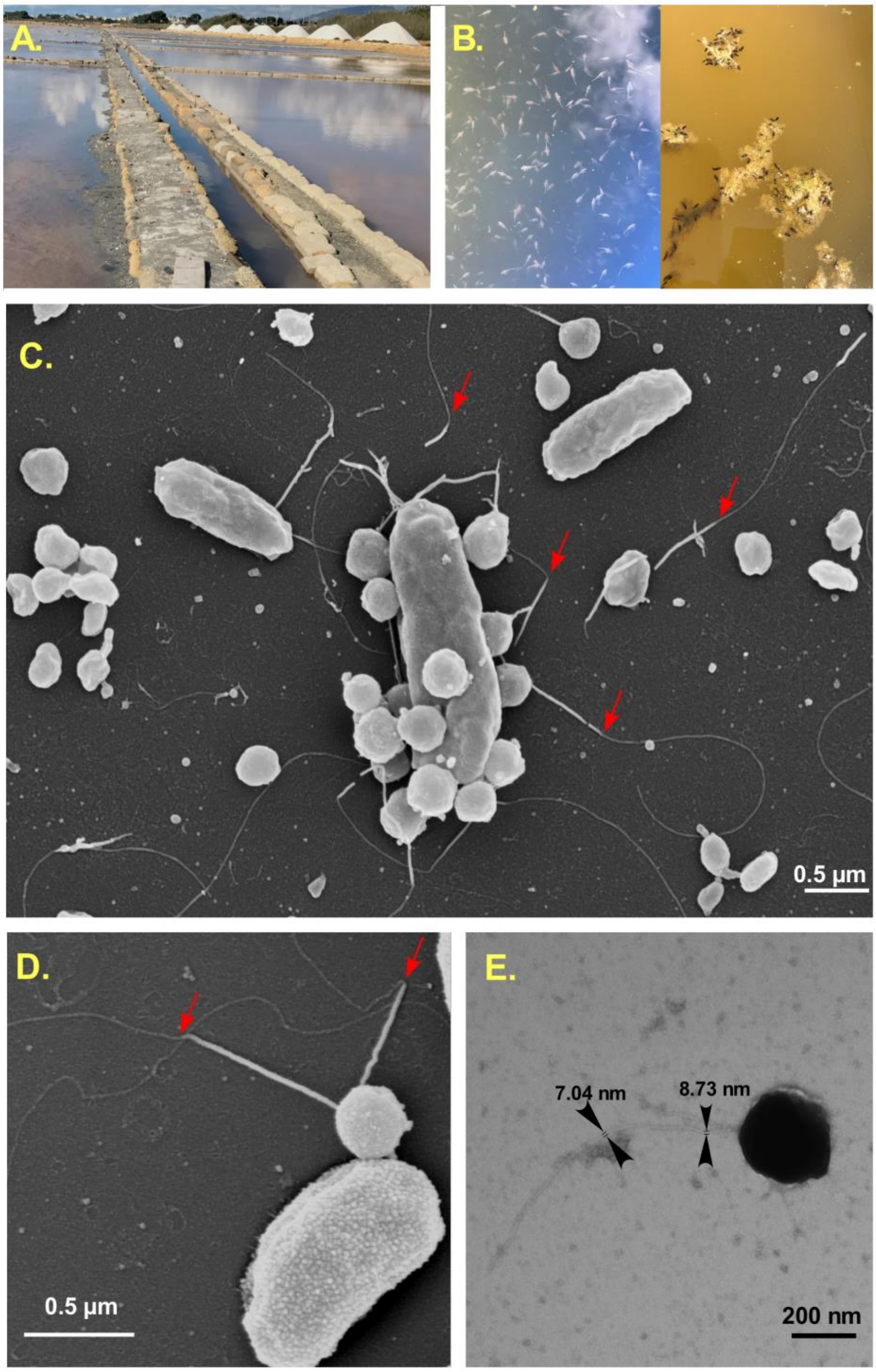


**Supplementary Figure 1| Source environment for Nanohaloarchaeota cultivation and electron micrographs of *Ca.* Nanohalobium constans LC1Nh in co-culture with its chitinolytic host, extreme halophilic *Halomicrobium* sp. LC1Hm.**

**A,** *Saline della Laguna* solar saltern system (37°51'48.70"N; 12°29'02.74"E), one of the ancient sites of salt production near Trapani (Sicily, Italy). **B,** Abundant presence of indigenous brine shrimps and brine flies was observed at the time of sampling. **C,** Field emission scanning electron microscopic (FESEM) images of the close interaction of nanohaloarchaeal tiny coccoid cells and its chitinotrophic host *Halomicrobium* sp. LC1Hm (elongated rods). We visualized that from one to up to 17 nanohaloarchael cells can be physically associated with the host cell. **C-E,** FESEM and TEM images of *Ca.* Nanohalobium constans cells possessing the pili-like structures as thick and long protein stalks of the archaella, which can unwind to thin filaments at points, evidenced by red arrows.

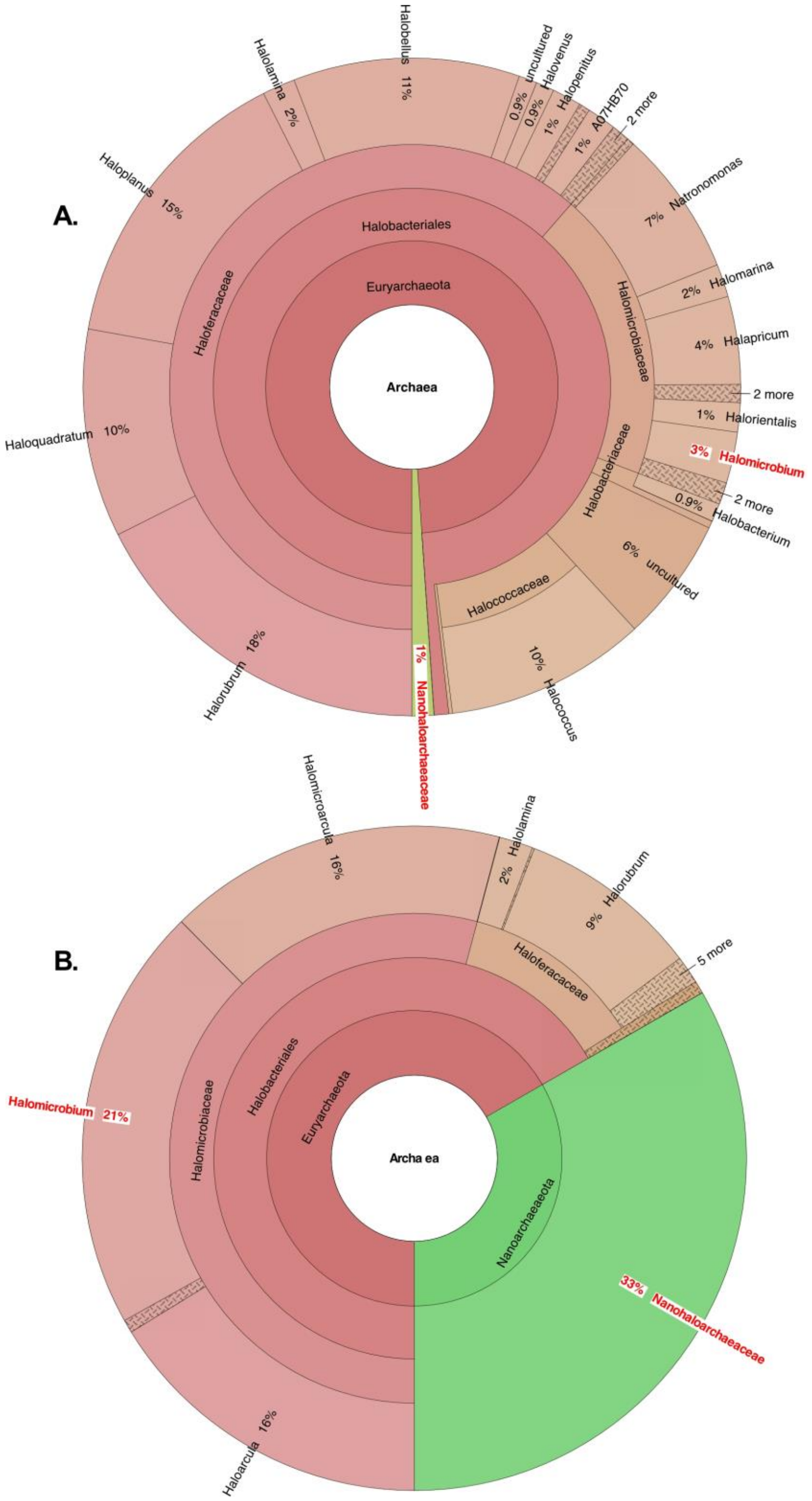


**Supplementary Figure 2| Taxonomic profiling of the archaeal communities found in crystallizer pond of Saline della Laguna (a) and in the chitin enrichment (b), based on 16S rRNA gene amplicons (sequenced by Illumina MiSeq platform).** 16S rRNA gene sequences were classified using the RDP classifier. For the taxonomic profile, only assignments with a confidence level of 0.8 were considered. The profile was visualised using Phylosift software and displayed with Krona tools. Nanohaloarchaea- and *Halomicrobium*-related fractions of amplicons are highlighted in red.

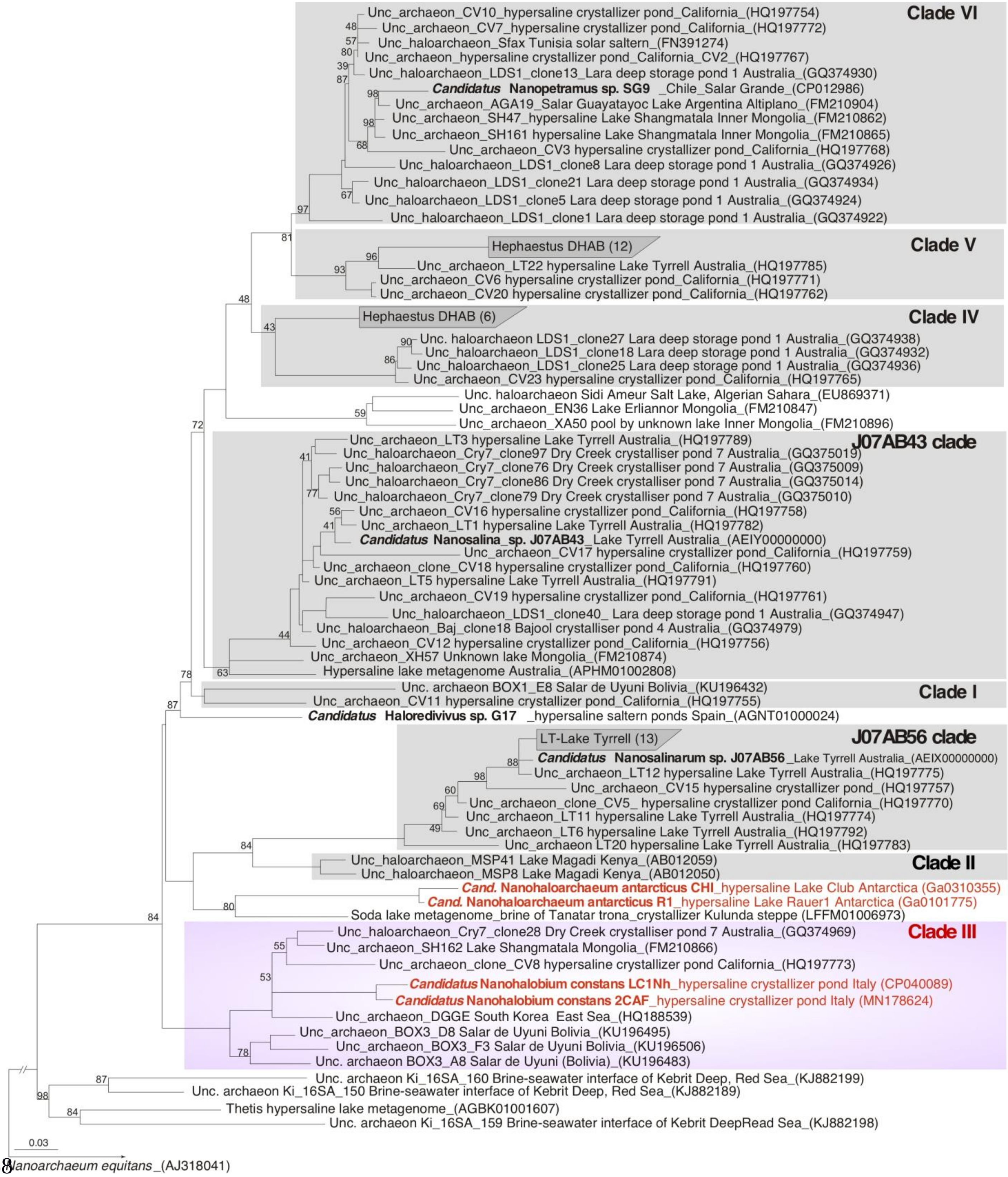


**Supplementary Figure 3| Phylogenetic position of *Candidatus* Nanohalobium constans LC1Nh within the *Nanohaloarchaeota*.** The tree was constructed based on alignment of the LC1Nh 16S rRNA gene sequence to the SILVA Release 132SSURef NR99 using phylogenetic inference under maximum parsimony criteria within the ARB software environment. The 16S RNA gene of *Nanoarchaeum equitans* (AJ318041) was used as an outgroup. Numbers at nodes represent bootstrap values (1000 replications of the original dataset). Values less than 40% were omitted. The scale bar represents the average number of substitutions per site. Cultivated nanohaloarchaea are highlighted in red.

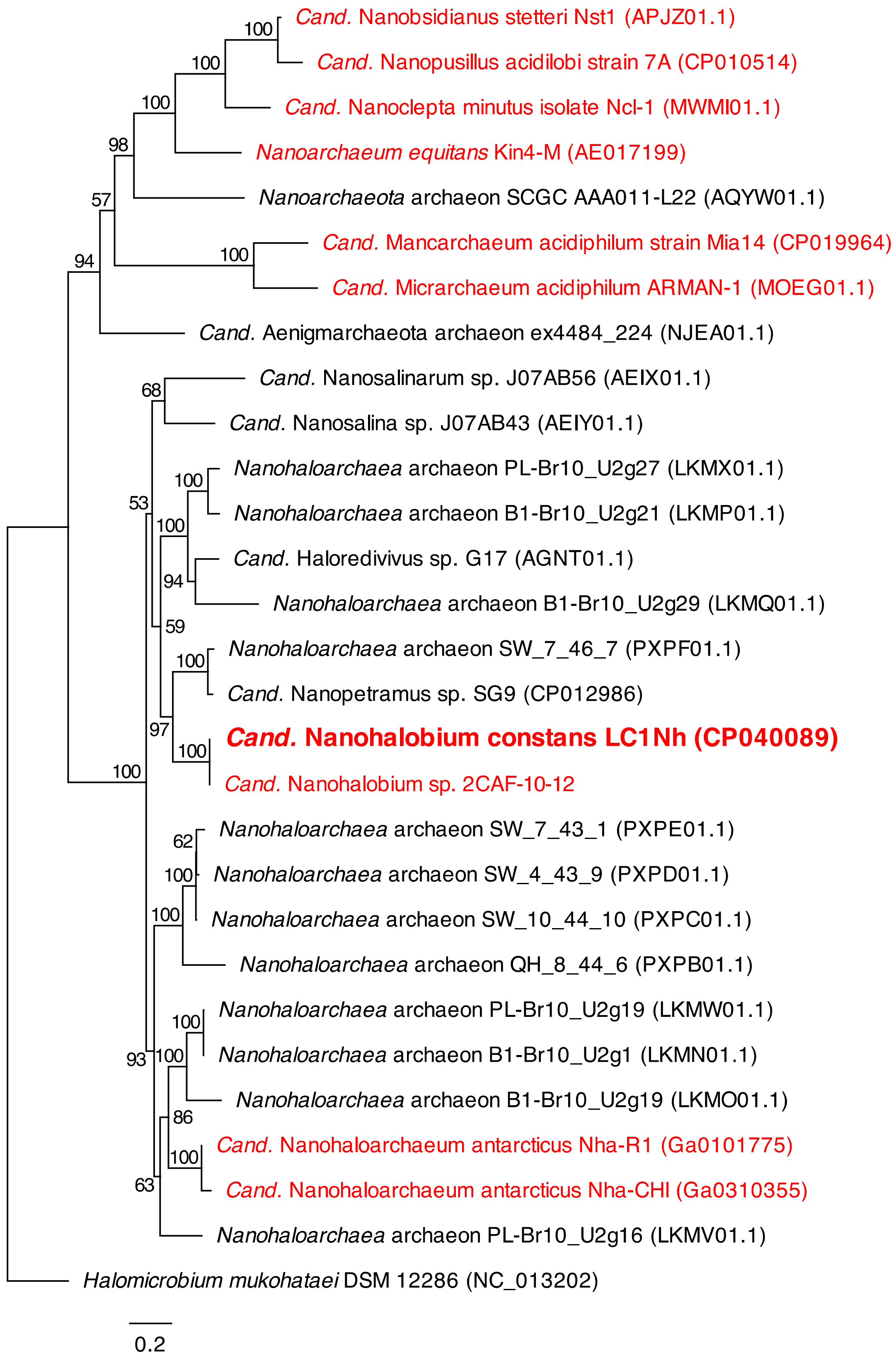


**Supplementary Figure 4| Maximum likelihood phylogeny of selected DPANN archaea based on the concatenated alignment of 11 ribosomal proteins**. The amino acids sequences were aligned by Clustal W 2.1 program with Blosum cost matrix, and the phylogeny was inferred by PhyML 3.0 plugin software inside Geneious 7.1, with Blosum62 substitution model and 1,000 bootstrap replicates. *Halomicrobium mukohataei* DSM 12286 was included as an outgroup. Bootstrap support values (if >40) are indicated for selected groups at the nodes. Cultured ectosymbiotic DPANN archaea are highlighted in red. The scale bar represents the average number of substitutions per site.

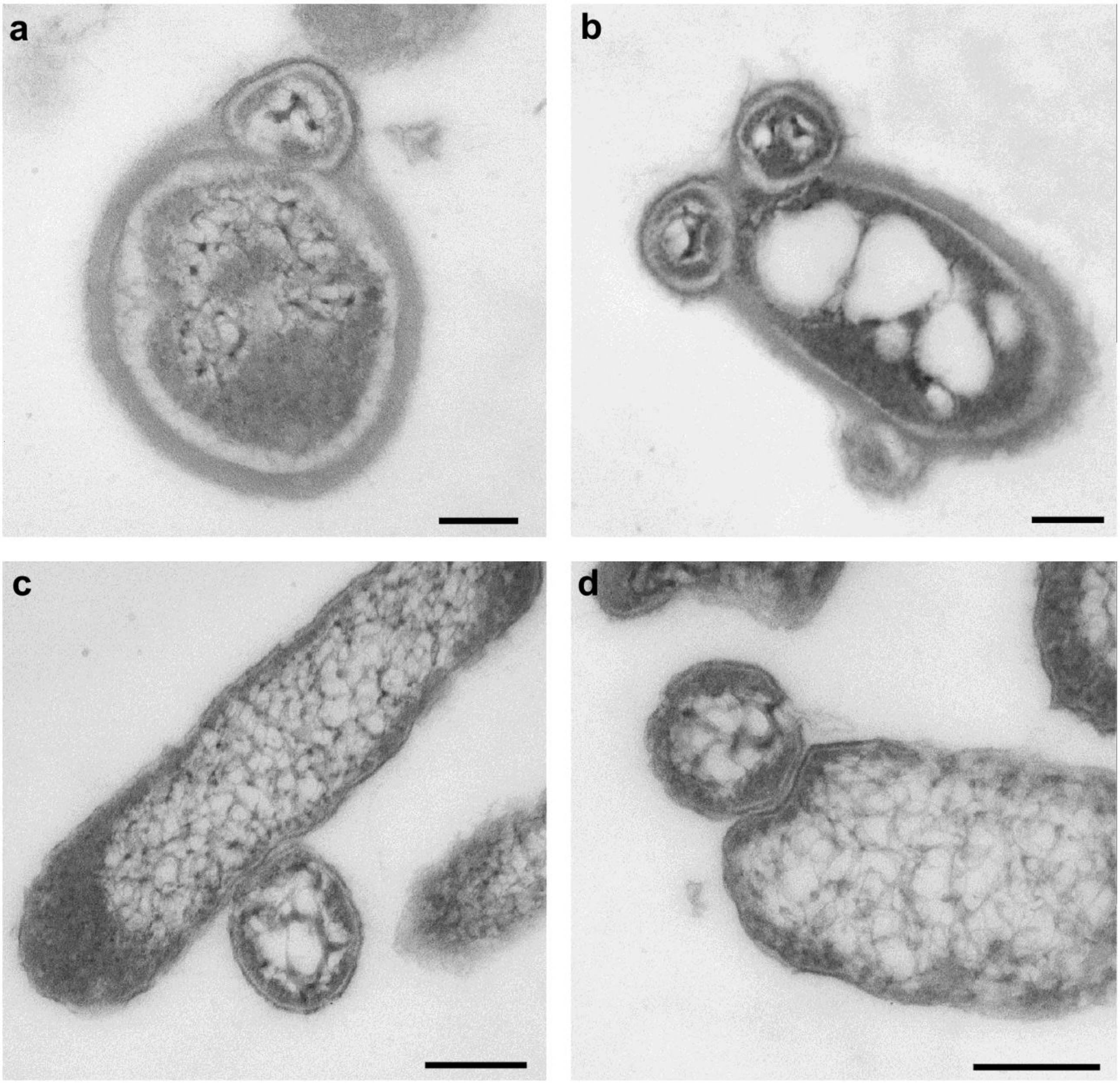


**Supplementary Figure 5|** TEM microscopic images demonstrate the intimate contact between *Ca.* Nanohalobium constans LC1Nh and the host *Halomicrobium* sp. LC1Hm cells. Growth on chitin (**a, b**) of *Halomicrobium* sp. LC1Hm is associated with the appearance of a thick electron dense external layer, which was not observed during tested growth on either monosaccharides (N-acetylglucosamine and dextrose) or disaccharides (maltose and cellobiose) (**c, d**). Scale bars represent 500 nm.

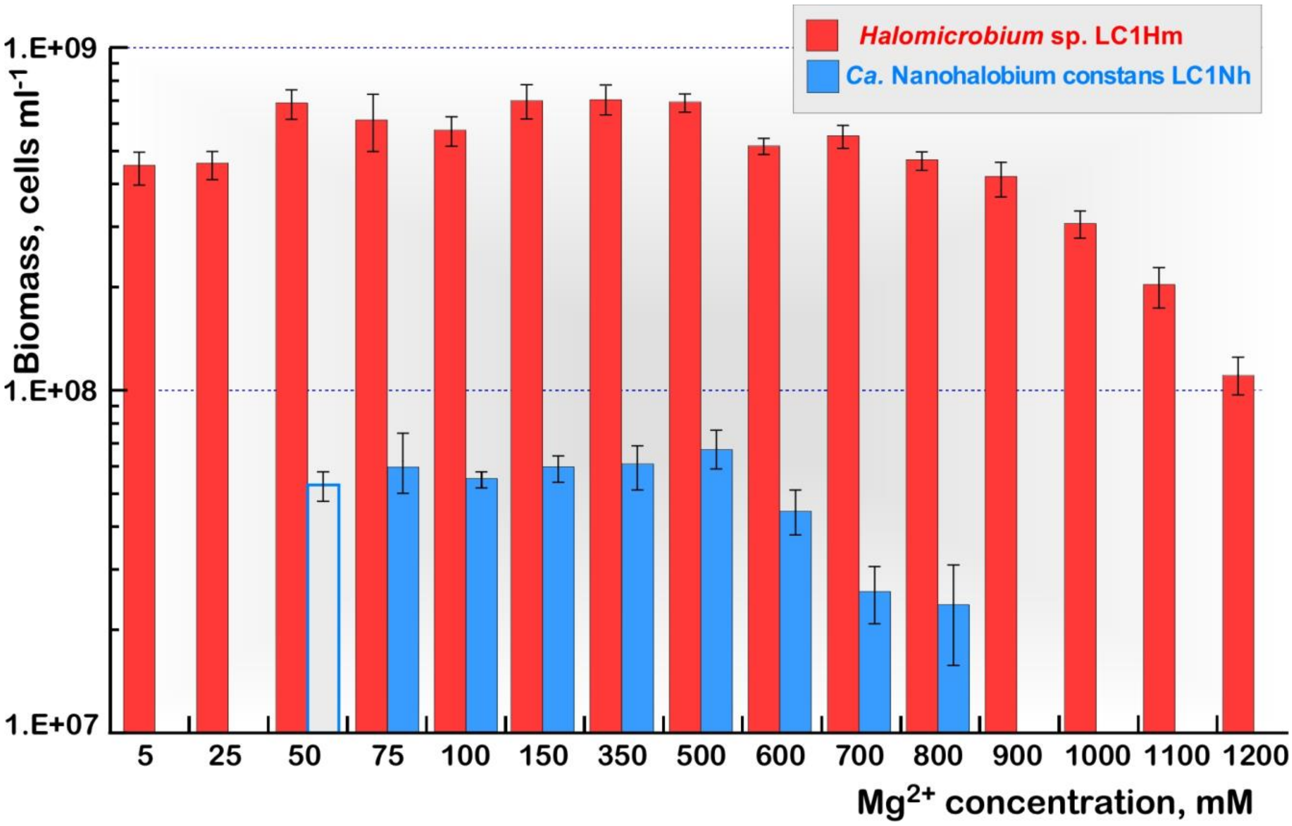


**Supplementary Figure 6|** Magnesium tolerance and magnesium dependence of extremely halophilic co-culture of *Halomicrobium* sp. LC1Hm and *Ca.* Nanohalobium constans LC1Nh. Cell densities was measured by qPCR analysis after ten days of cultivation under microaerophilic conditions at 40°C and pH 7.2 using chitin (5 g $l^{-1}$) as the sole carbon and energy source. Total salinity of LC culture medium was maintained as 240 g $l^{-1}$ by varying either sodium or magnesium salts. Growth of *Ca.* Nanohalobium constans LC1Nh at 50 mmol $Mg^{2}$+ concentration, shown as grey column, was observed only in the first generation of the LC1Hm+LC1Nh co-culture, originally grown at 315 mM $Mg^{2+}$. Error bars (standard deviations) are based on three culture replicates.

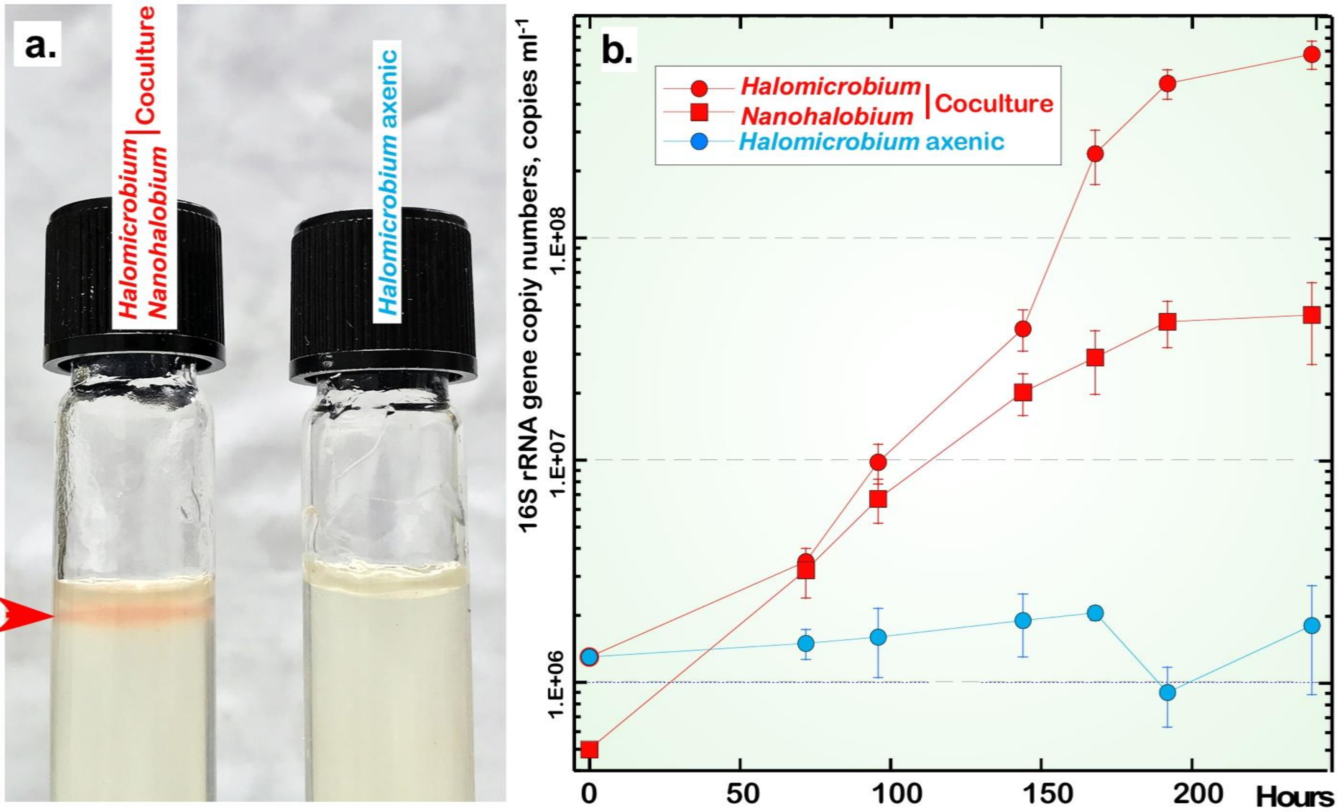


**Supplementary Figure 7|** Growth on glycogen (2 g $l^{-1}$) of *Halomicrobium* sp. LC1Hm in co-culture with *Ca.* Nanohalobium constans LC1Nh. (**A**.) Microaerophilic growth in solidized (1.5% of agar, w/w) LC medium. Red arrow points the zone of microaerophilic growth. Gradient of oxygen was achieved by placing of 1 ml of 2 mM $Na_2S$ on the bottom of each Hungate tube. (**B**.) Time course of growth of *Halomicrobium* sp. LC1Hm in pure (axenic) culture and in co-culture with *Ca.* Nanohalobium constans LC1Nh. Error bars (standard deviations) are based on three culture replicates.

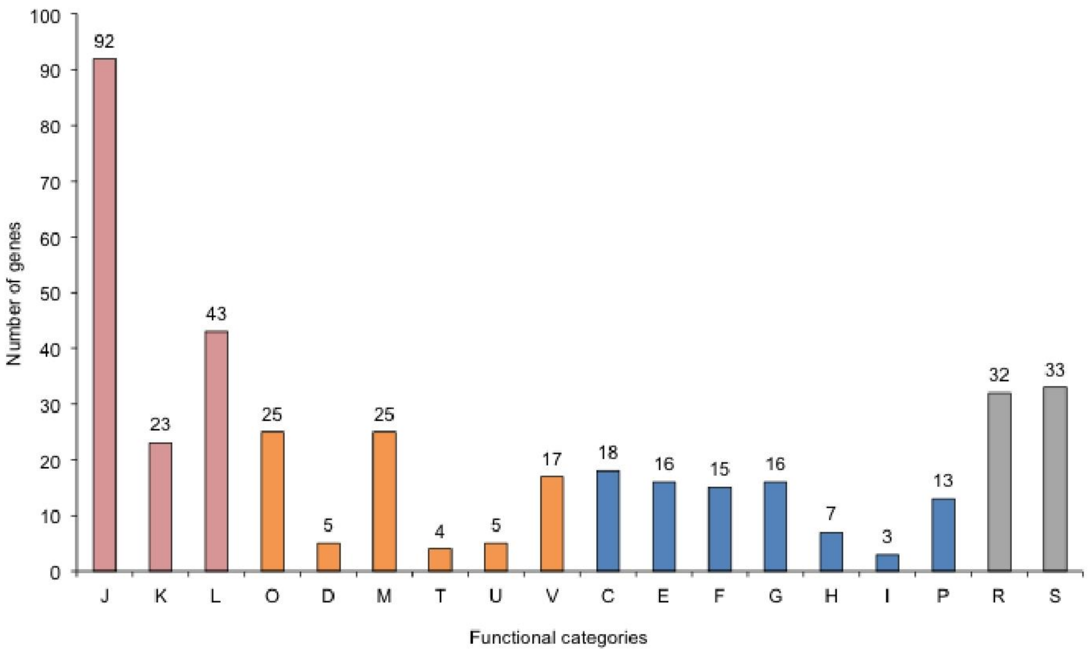


**Supplementary Figure 8| Gene functional categories of *Ca.* Nanohalobium constans LC1Nh.** Functional categories are designated according to the NCBI COG resource as follows. Shown in pink: J, translation, ribosomal structure and biogenesis; K, transcription; L, replication, recombination and repair; shown in orange: O, posttranslational modification, protein turnover, chaperones; D, cell cycle control, cell division, chromosome partitioning; M, cell wall/membrane/envelope biogenesis; T, signal transduction mechanisms; U, intracellular trafficking, secretion, and vesicular transport; V, defence mechanisms; shown in blue: C, energy production and conversion; E, amino acid transport and metabolism; F, nucleotide transport and metabolism; G, carbohydrate transport and metabolism; H, coenzyme transport and metabolism; I, lipid transport and metabolism; P, inorganic ion transport and metabolism; shown in grey: R, general function prediction; S, function unknown.

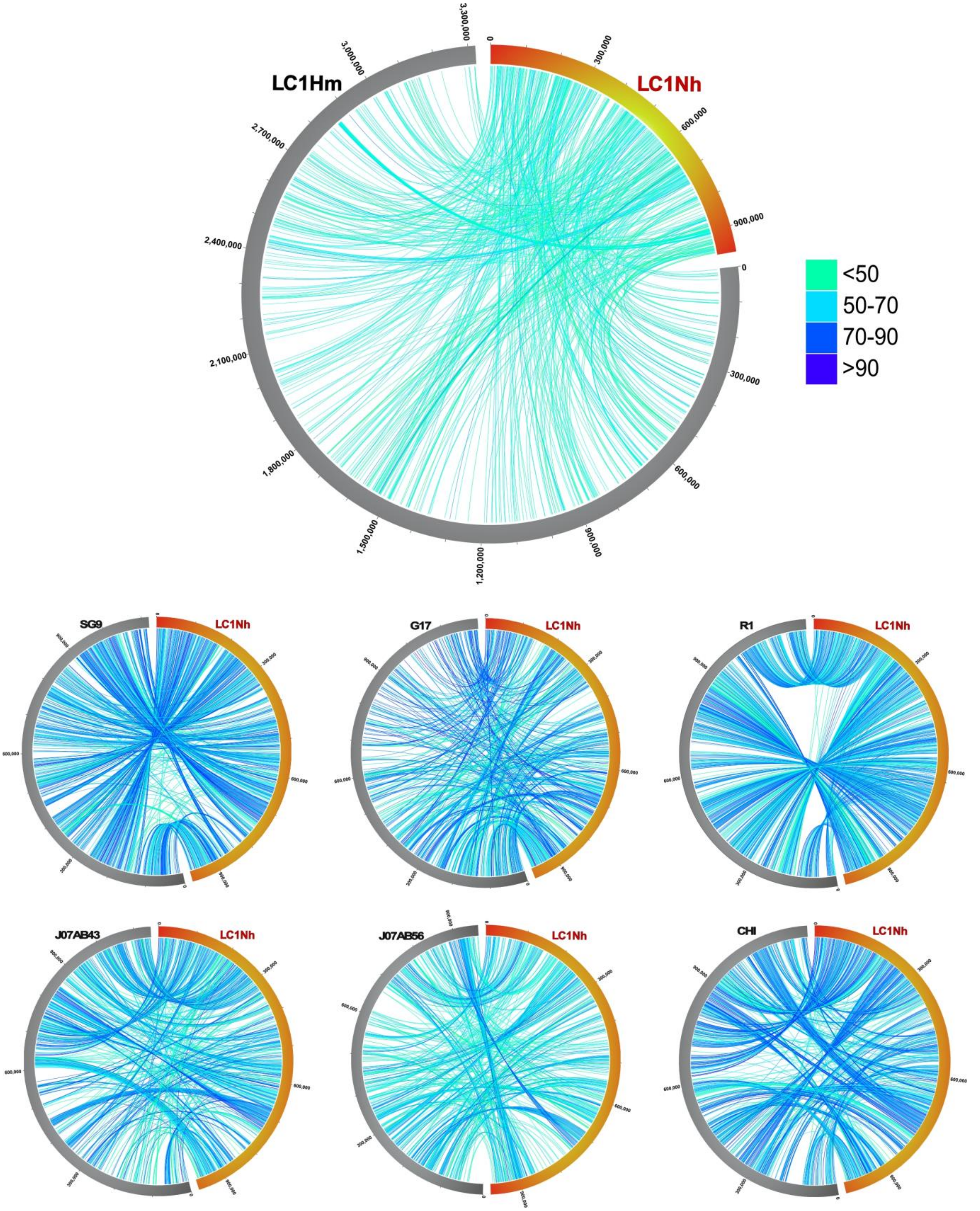


**Supplementary Figure 9|** Circos-based genome alignments of *Ca.* Nanohalobium constans **LC1Nh** versus the host *Halomicrobium* sp. **LC1Hm** and six nanohaloarchaea: *Ca.* Nanopetramus sp. **SG9**, *Ca.* Haloredivivus sp. **G17**, *Ca.* Nanosalina sp. **J07AB43**, Ca. Nanosalinarum sp. **J07AB56**, *Ca*. Nanohaloarchaeum antarcticus **R1** and *Ca*. Nanohaloarchaeum antarcticus **CHI**. Links indicate pairs of orthologous genes between the genomes, the colour is scaled to percentage of amino-acid identity levels.

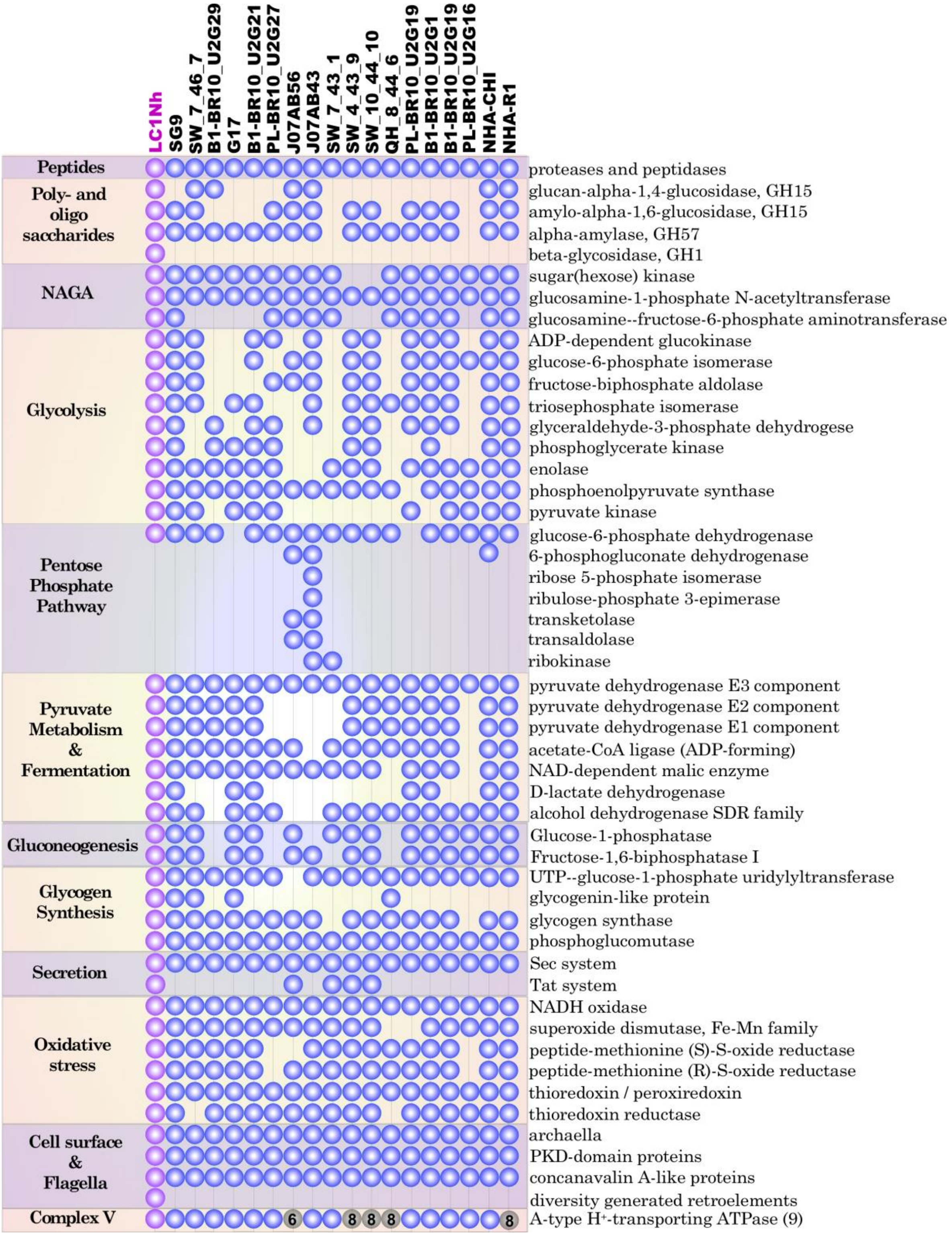


**Supplementary Figure 10| Comparative metabolic analysis of the 19 nanohaloarchaeal genomes.** Proteins of central carbon metabolism and proteins putatively involved in electron transfer, oxidative stress, secretion, extracellular matrix, archaella and diversity-generating retroelements. *Ca.* Nanohalobium constans LC1Nh proteins are shown in magenta. The A-type $H^+$ translocating ATPase complex (9 subunits) is intact in 13 nanohaloarchaeal genomes, whereas incomplete complexes are shown by grey colour with the numbers of subunits found. The list is not mutually exclusive as a given protein can have more than one function or domain and was counted in each appropriate category.

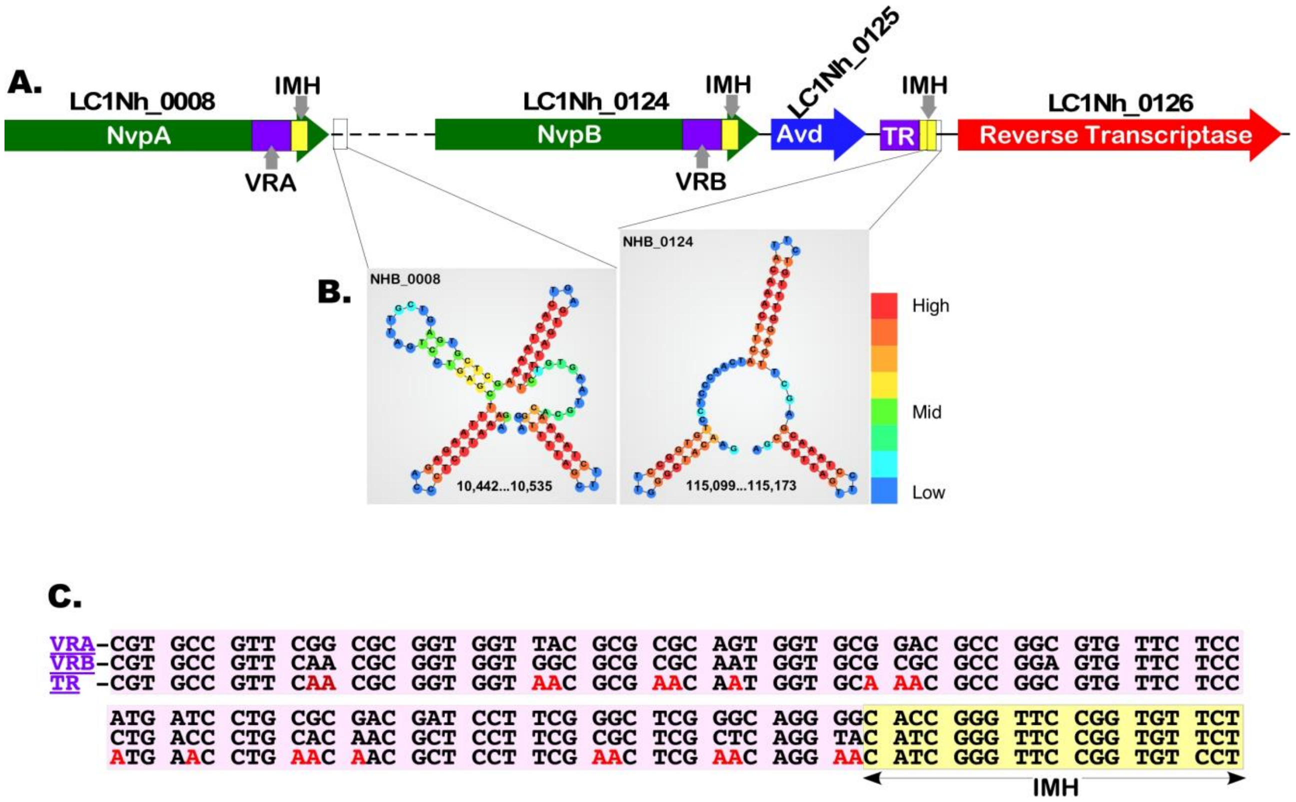


**Supplementary Fig. 11| Diversity-generating retroelements (DGR) locus in *Ca.* Nanohalobium constans LC1Nh genome.**

(**A.**) DGR locus consists of consists of the accessory variability determinant (Avd, LC1Nh_0125) and the error-prone reverse transcriptase (RT, LC1Nh_0126). Between the Avd and RT genes the LC1Nh genome contains a 95-bp long template region (TR). TR is similar to the variable regions (VRA and VRB) of two proteins, LC1Nh_0008 and LC1Nh_0123 (80% and 84%, respectively). At the 3'-end of the VRA, VRB and TR regions, the LC1Nh DGRs system composes of three identical 19 bp-long sequences, coined as initiation of mutagenic homing sequences (IMH). (**B.**) Two hairpin/cruciform structures downstream of the VRA and VRB were evident in the LC1Nh genome. (**C.**) Alignment of VRA and VRB regions of Nanohalobium variable proteins NvpA and NvpB (LC1Nh_0008, 0123) with TR.

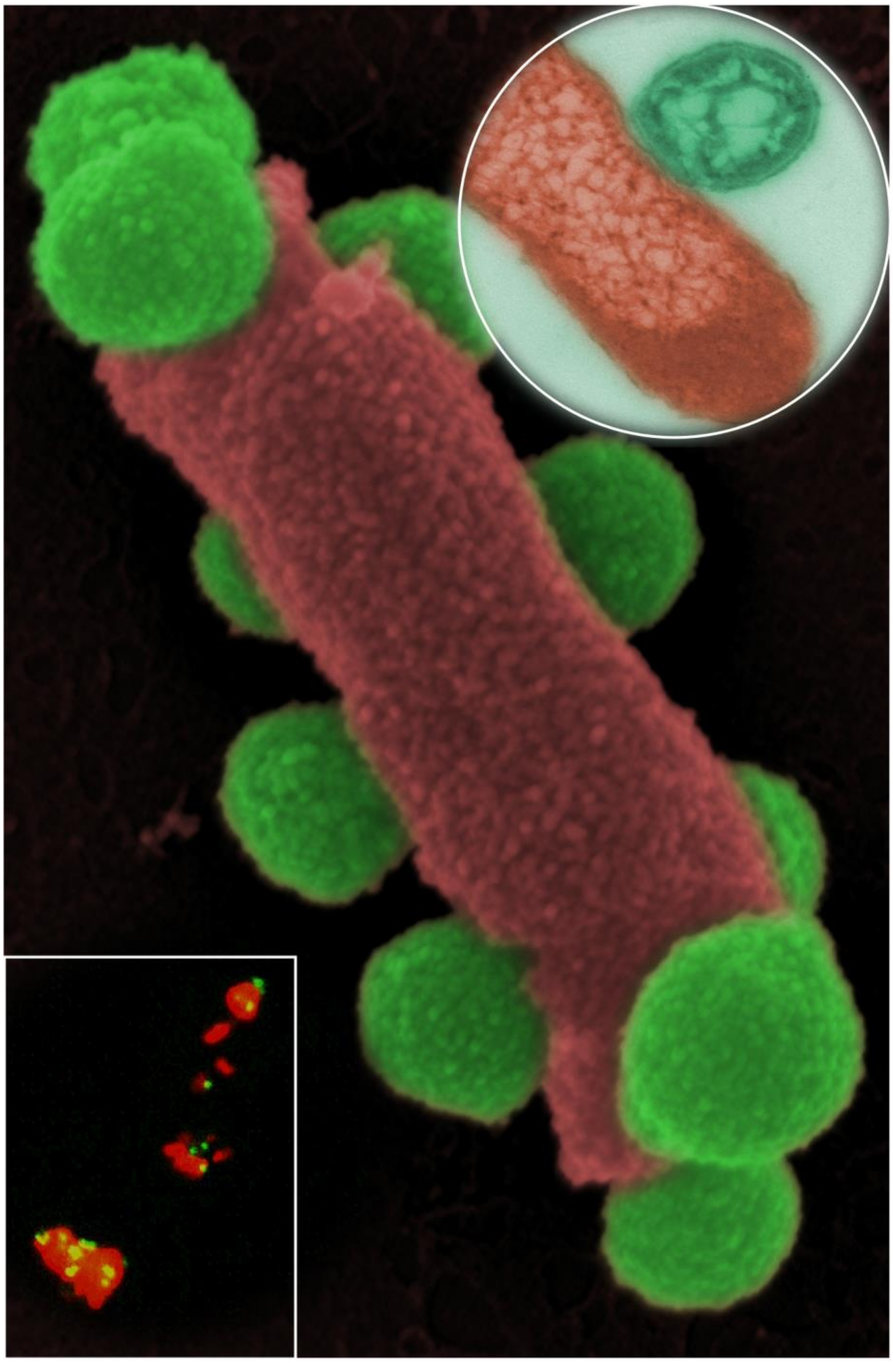

**Fig. 12: Potential cover caption. Could the association between ectosymbiotic nanohaloarchaeon and chitinolytic haloarchaea be beneficial to the host?**

The collage combines a CARD-FISH fluorescence microscopic image (rectangular insert in lower left side) with FESEN scanning (central image) and transmission microscopic image of an ultrathin section (circular insert in upper right part) to demonstrate the intimate contact between Nanohaloarchaeota and their host cells. The image was created with Adobe Photoshop 5.0 by colorizing scanning and transmission electron microscopic images, according to the colours of CARD-FISH fluorescence. The fluorescence microscopic image was adjusted in contrast and brightness.
